# Evaluation of Multiplexed Liquid Glycan Array (LiGA) for Serological Detection of Glycan-binding Antibodies

**DOI:** 10.1101/2025.06.23.660925

**Authors:** Revathi Reddy, Eric Carpenter, Anne Halpin, Mirat Sojitra, Chuanhao Peng, Guilherme Meira Lima, Xiaochao Xue, Kejia Yan, Jean Pearcy, Maria Ellis, Bruce Motyka, Todd L. Lowary, Lori West, Ratmir Derda

**Affiliations:** Department of Chemistry, University of Alberta, Edmonton, AB T6G 2G2, Canada; Department of Pediatrics, University of Alberta, Edmonton, AB, Canada; Alberta Transplant Institute and Canadian Donation and Transplantation Research Program, University of Alberta, Edmonton, AB, Canada; Department of Laboratory Medicine and Pathology, University of Alberta, Edmonton, AB, Canada; Institute of Biological Chemistry, Academia Sinica, Taipei Taiwan; Institute of Biochemical Sciences, National Taiwan University, Taipei Taiwan; Department of Surgery, University of Alberta, Edmonton, AB, Canada; Department of Medical Microbiology and Immunology, University of Alberta, Edmonton, AB, Canada

## Abstract

We test the performance of the multiplexed liquid glycan array (LiGA) technology in serological assays. Specifically, we use LiGA to detect ABO blood group antibodies in human serum. This LiGA, which we name ABO-LiGA, contains ABO blood group trisaccharide glycans with an ethylazido aglycone conjugated to groups of ten multi-barcoded M13 particles carrying dibenzocyclooctyne (DBCO) on p8 proteins. ELISA clonal binding assays to anti-A/B antibodies confirmed the functional performance of ABO-clones and aligned with next-generation sequencing (NGS) of the mixed clones. Multiple DNA-barcoded technical replicates in LiGA allow for quantification of reproducibility and robustness as determined by the Z’-score using NGS. We then tested ABO-LiGA for specific detection of IgG and IgM anti-A and anti-B IgG and IgM antibodies in human serum samples. Comparison of antibody binding responses in sera from 31 healthy donors to ABO-LiGA with an ABO-Luminex-based method revealed consistent responses to LiGA-ABO but also minor deficiencies of ABO-LiGA such as low robustness of the current assay format and a limited ability to detect low intensity antibody responses. Some results point to undesired interactions of serum antibodies with small-footprint glycans conjugated to phage via the bulky DBCO moiety. This report illuminates the path for future development of LiGA-based serological assays and suggests the need to develop alternative methods for conjugating glycans to phage to avoid liabilities of the hydrophobic DBCO moiety.

## Introduction

Glycans coat the surfaces of every cell across all domains of life (Smith and Bertozzi 2021). Their prominent display on cell surfaces is linked to the central role of glycans in glycan–antibody interactions in the present-day immune system and peptidoglycan-based microbe-associated molecular patterns (MAMPs) (Johannssen and Lepenies 2017; Temme et al. 2021). Binding of immunity lectins and antibodies, components of the innate and adaptive immune systems, to glycans plays a critical role in self versus non-self-discrimination by the immune system. The collection of glycans across all living things is incredibly diverse and often taxonomically gated. Glycan recognition profiles of antibody repertoires can, in theory, be used to analyze the glycans and species to which an organism has been exposed (Marglous et al. 2024). Antibody–glycan recognition analysis in serum is critical for development of tool antibodies and therapeutic antibodies (McKitrick et al. 2020). Analysis of antibody–glycan interactions is also important to understand the fundamental plasticity displayed by anti-carbohydrate antibodies in their recognition of carbohydrates (Kappler and Hennet 2020). These collective investigations are possible using high-throughput systems for measuring interactions between diverse carbohydrates and diverse antibodies in human serum (Marglous et al. 2024). In this report, we test whether recently a reported DNA-encoded, multivalent display of glycans on bacteriophages (Sojitra et al. 2021) can be used to detect glycan-binding antibodies in human serum.

Glycan arrays are an established high-throughput tool that allows characterization of the interactions between glycans and glycan binding receptors (Blixt et al. 2004; Geissner et al. 2014; Oyelaran and Gildersleeve 2009; Xia and Gildersleeve 2015). Glycan arrays confirmed that human serum has a diverse assortment of carbohydrate binding antibodies generated in response to glycans present on the cell surface of pathogens, tumor cells and vaccines (Geissner et al. 2014; Kappler and Hennet 2020; Marglous et al. 2024). Precise control of multivalent presentation is one drawback of conventional “solid” printed arrays in which glycans are immobilized on a glass surface leading to significant variability (Temme et al. 2019). Serial dilution has been used to produce different epitope densities, but characterization of such densities on glass slides is not trivial and often impossible. This problem can be minimized by preparing neoglycoproteins of definitive density (Zhang and Gildersleeve 2011); printing such glycoproteins on the surface of the arrays made it possible to detect populations of glycan-binding proteins and antibodies that were not possible to resolve with traditional arrays (Oyelaran et al. 2009). Similar approaches exist in which prospectively manufactured multivalent constructs are printed on the surface (Narla et al. 2015). We reasoned that a controlled multivalent display of glycans on bacteriophage virions (Sojitra et al. 2021) may offer a convenient platform for detection of anti-glycan antibodies in human serum.

A limitation of conventional slide-based glycan arrays is the necessity of specialized array printing and array reader instruments that are not widely available, particularly in clinical laboratories (Anne Halpin et al. 2024). Therefore, bead-based arrays (Purohit et al. 2018) have been developed to allow obtaining array-like information from flow cytometer-like system readily available in clinical laboratories (A. Halpin et al. 2018; A. Halpin et al. 2020; A. M. Halpin et al. 2025). Glycan arrays were inspired by DNA microarrays (Blixt et al. 2004); however most questions that required DNA-arrays are now answered by next-generation sequencing (NGS). A shrinking market share for DNA microarrays may eventually lead to discontinuation of the hardware for printing and reading microarrays. In the past five years, multiple reports showcased that classical glycobiology tools can be transitioned from an array-based format to a DNA-encoded format. Examples include DNA-encoded libraries of glycans conjugated to DNA strands (Kondengaden et al. 2020; Thomas et al. 2017; Yan et al. 2019), collections of lectins tagged by DNA directly (Minoshima et al. 2021; Odaka et al. 2022) or indirectly (Kearney et al. 2021; Lima et al. 2024), multivalent display of N-linked glycans displayed on and encoded by RNA

(Horiya et al. 2017) and enabling technologies such as enzymatic assembly of monosaccharide-modified DNA (Kromer et al. 2024) and on-DNA synthesis of glycoconjugates (Ling et al. 2023; Zhao et al. 2023). We demonstrated that a multivalent, DNA-encoded “liquid” glycan array (LiGA) of glycans displayed on bacteriophage can use NGS to obtain glycan-array-like outcomes (Sojitra et al. 2021) and profiling of glycan-protein interactions on the surface of live cells *in vitro* (Sojitra et al. 2025) and *in vivo* (Lin et al. 2023). LiGA is built on prospective silent DNA barcoding (SDB) of M13 virions (Tjhung et al. 2016). These SDBs have been used extensively to encode chemical modifications to phage-displayed libraries (Chou et al. 2018; Ekanayake et al. 2021; Lima et al. 2022; Ng et al. 2018; Sojitra et al. 2021; Triana and Derda 2017; Vinals et al. 2019; Wong et al. 2021). The LiGA employs M13 virions with an SDB in the genome to encode identity and density of glycans displayed on phage. 2700 copies of outer phage coat protein allow for multivalent display of glycans (between 10–1500 copies per virion). The DNA barcodes inside the phage, with a theoretical diversity of ∼12 billion variants (Sojitra et al. 2021), can encode a vast number of variables such as composition and density of glycans to provide information that can decode the interaction of glycans with proteins and cell-based targets. In this manuscript, we draw inspiration from report by Casey Krusemark, Emily Dykhuizen and co-workers (Denton et al. 2018) to encode multiple technical replicates in the same solution by DNA; this multi-SBD (MSDB) concept is used in this manuscript to test robustness of LiGA technology in heterogeneous serum samples.

An ideal platform for serological tests and detection of glycan binding antibodies in human serum should have the following properties: (i) Reading by instruments already available in the clinic such as flow cytometers or DNA sequencers. (ii) A simple standardized workflow and built-in quality control procedures (QC). (iii) Ability to display and test glycans with different linkages and densities. The LiGA platform (Sojitra et al. 2021) has the potential to fulfill these requirements. In this publication, we perform a preliminary test of the capacity of LiGA to detect antibodies against carbohydrates in the complex environment of human serum, a media that contains billions (Fanning et al. 1996) of diverse antibodies. To test the potential of LiGA to measure the enrichment of specific glycans by specific antibodies from a mixture of glycans, we employed DNA barcoding to encode replicates of identical glycoconjugates. Such encoding effectively assesses multiple technical replicates within the same bulk assay and quantifies robustness and reproducibility of antibody–glycan interactions. This publication is focused entirely on detection of ABO blood group antibodies using ABO-LiGA. Other anti-glycan antibodies in serum (Bovin et al. 2012; Kappler and Hennet 2020; S. M. Muthana and Gildersleeve 2016a; Pochechueva et al. 2016; Temme et al. 2022), in principle, can be analyzed by LiGA as well; however, we defer these experiments to subsequent reports.

LiGA assay is wholly dependent on Illumina “sequencing by synthesis”. It therefore may be impacted by standard error types: “insertion” and “deletion” of a nucleotide as well as a “substitution” error or “termination”. We employ paired-end sequencing for detecting and correcting some of these errors. Negative binomial distribution (Buller et al. 2009), Poisson distribution (Kuai et al. 2018; MacConnell and Paegel 2017), the Z’ score metric (Denton et al. 2018; Faver et al. 2019) and machine learning approaches (Amigo et al. 2018; Komar and Kalinic 2020) have been used to quantify the biases in NGS. However, many complex error correction methods can be avoided because LiGA uses a fixed dictionary of silent barcodes. A predetermined Hamming (Hamming 1950) distance (H) between all barcodes allows their identification even if the sequencing procedure introduces random errors; discussion of such error correction can be found elsewhere (Buschmann and Bystrykh 2013; Bystrykh 2012; Hamming 1950; Levenshtein 1966). In short, despite PCR or Illumina NGS errors, a H>3 distance between all barcodes makes it possible to recognize barcodes after most simple “typos” introduced by the sequencing procedure. The error correction is simplified owing to the fixed dictionary of barcodes as well as the fixed identify of forward and reverse priming sites (Matochko and Derda 2013). After all error corrections, we use the Z’ score to evaluate the robustness of the LiGA-based approach in serological assays. This allows us to uncover evidence for barcode outliers that might skew assay outcomes as well as identify systematic biases that we attribute to intrinsic liabilities of the strategy used for conjugation of glycans to the phage particle.

## Results

The preparation of ABO-LiGA (**Figure 1a-c**) employed previously-reported strain promoted azide– alkyne cycloaddition (SPAAC) between dibenzocyclooctane (DBCO) ligated to the pVIII protein of the M13 virion and azido glycans (Sojitra et al. 2021). Prior to each conjugation, we mixed ten clones, each with a unique SDB, to create a multiplexed SDB (MSDB) (**Figure 1c, Figure S1-S2**). This procedure associated a specific glycan with ten DNA clones (**Figure S3**). As a result, a single measurement by the LiGA assay yielded 10 replicates of the binding response for the same glycan in the same assay. Typical four independent binding experiments, hence, give giving rise to 4×10 data points per glycan, per experiment. To each MSDB-barcoded phage, we conjugated Tri-AN3 (A blood group trisaccharide), Tri-BN3 (B blood group trisaccharide) or Di-N3 (O blood group disaccharide), each with an ethyl-azido aglycone linker (**Figure 1d, Figure S4**). These three glycans are referred to as ABO-glycans in the remaining manuscript.

**Figure 1:**
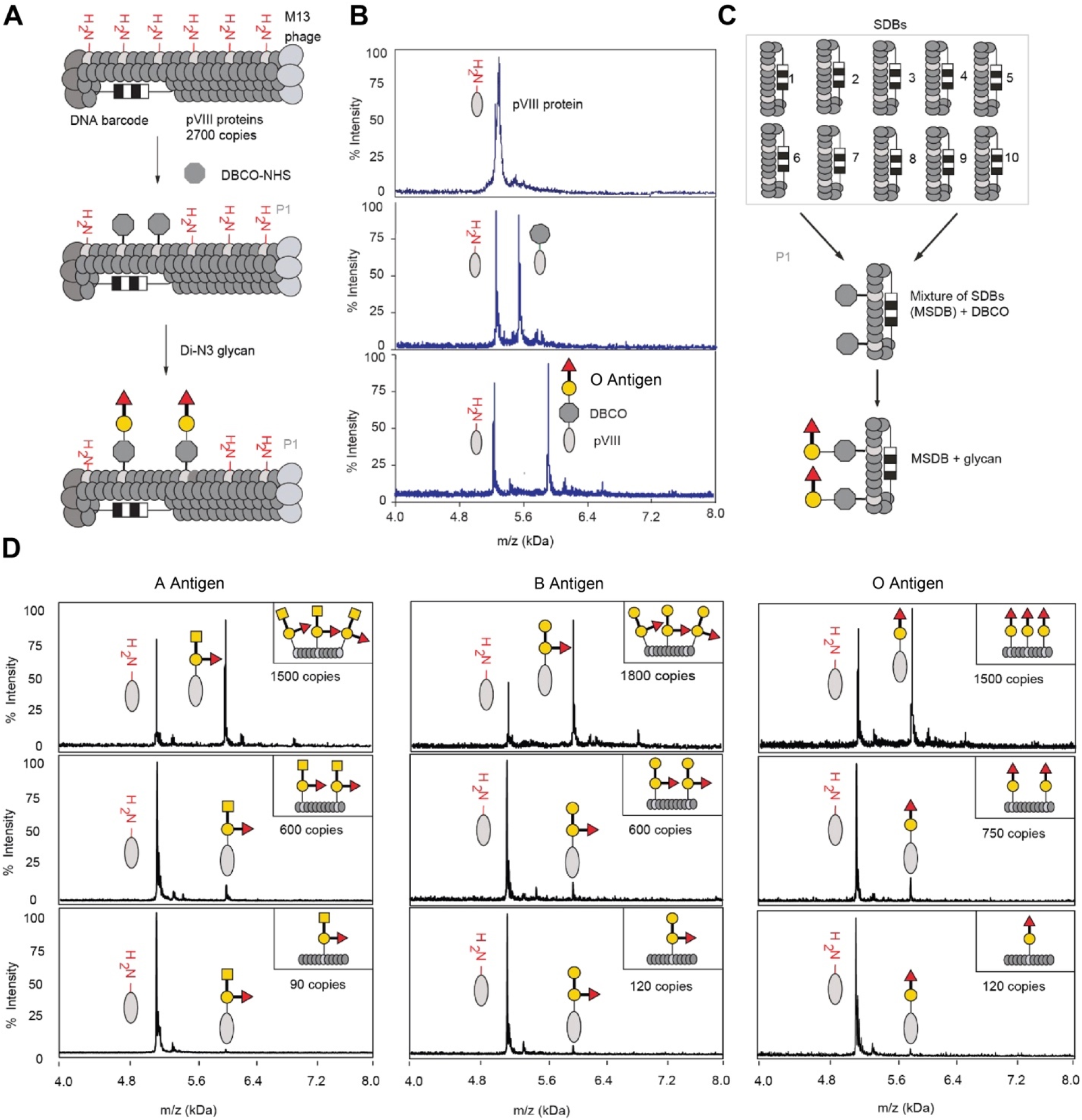
Synthesis and characterization of ABO phage glycoconjugates. **a)** Representation of the two-step chemical glycosylation of phage with the H disaccharide antigen. **b)** MALDI mass spectrometry characterization of the starting material (protein pVIII), the alkyne-functionalized product (DBCO–pVIII, P1) and the glycoconjugate product (P2). **c)** Representation of the synthesis of MSDB glycan conjugates. **d)** MALDI-TOF spectra of MSDB conjugated to ABO glycans with varying densities. This modification density was achieved by controlling the reaction conditions and concentration of DBCO-NHS ester. For example, by changing the concentration of DBCO-NHS from 1.5 mM to 1 mM and 0.5 mM, we could produce conjugates Tri-AN3-[1500], Tri-AN3-[600], Tri-AN3-[90], Tri-BN3-[1800], Tri-BN3-[600], Tri-BN3-[120], Di-N3-[1500], Di-N3-[750], Di-N3-[120]. Numbers in square brackets indicate the number of glycans per phage particle.

We published that the relationship between binding of glycans to GBPs, and the density at which glycans are displayed on the surface of the M13 display, are non-linear and even bimodal. In the latter case, an increase in glycan density leads to a decrease in binding due to steric occlusion (Blixt et al. 2004; Dam and Brewer 2010; Oyelaran and Gildersleeve 2009; Sojitra et al. 2021). Glycan density impacts the binding to two binding sites on IgG (Mian et al. 1991) and ten binding sites on IgM subtypes (Oyelaran and Gildersleeve 2009); such avidity-based responses are known to accentuate affinity of antibodies towards glycans (Toone 2002). Based on a published procedure (Sojitra et al. 2021), we varied the concentration of dibenzocyclooctyne-*N*-hydroxysuccinimidyl ester (DBCO–NHS) to acylate a pre-determined number of pVIII proteins on M13 phage (**Figure 1d**). The peak intensities in MALDI semi-quantitatively represent the fraction of a total of 2700 copies of pVIII modified by DBCO (**Figure 1d**). SPAAC ligation with an excess (2□mM) of ABO glycans quantitatively converted all DBCO–pVIII to the corresponding glycoconjugate-pVIII. Disappearance of the pVIII–DBCO peak in the MALDI spectrum confirmed the complete consumption of DBCO. This procedure ligated ABO glycans at three different densities—low [90–120], medium [600–800] and high [1500–1800]—to test the role of glycan presentation. The numbers in square brackets indicate the number of glycan particles per phage particle and they provide estimate for the *average* spacing between glycans on the bacteriophage surface (Sojitra et al. 2021). The 1500–1800 density corresponds to ∼50% of the pVIII proteins carrying an ABO glycan with the average spacing between ABO glycans being similar to the spacing between the pVIII proteins (3–4 nM).

### Validation by Enzyme Linked Immunosorbent Assay and Plaque Forming Unit Assays

We repurposed the classical phage-ELISA (Barbas et al. 2004) to evaluate the functional integrity of our individual glycosylated phage constructs, and to optimize the coating density of anti-A and anti-B antibodies (**Figure 2a-c**). In binding to anti-A antibody coated wells, phage displaying the A-trisaccharide exhibited the highest signal while phage conjugated to the H-disaccharide exhibited 8–10-fold lower binding. Similarly, phage displaying the B-trisaccharide exhibited the highest binding with anti-B antibody and a 15–17-fold lower binding with disaccharide H. Both A and B phages showed significantly lower binding to plates coated with bovine serum albumin (BSA) (**Figure 2b**). For both anti-A and anti-B antibodies, an optimal signal-to-noise ratio was obtained at 1:1000 and 1:2000 dilutions. Titration of a glycophage on a surface coated by these two dilutions of antibodies yielded a dose response with EC_50_ in the 5–8×10^6^ PFU/well range for both anti-A and anti-B antibodies (**Figure 2c**). Given well volume (100 μL), this number corresponds to 5–8×10^10^ virion/L or a 0.5–0.6 nanomolar concentration. The EC_50_ of detection is many orders of magnitude below the K_d_ of typical glycan–antibody interactions, an observation that highlights the avidity arising from interactions between multiple glycans on the 700 nm-long phage and multiple antibodies on the surface. Based on ELISA from antibody dilution screens, antibodies were diluted 1:1000 to ensure that the wells have optimal coating and the input concentration of LiGA was calculated to contain at least 10^6^ copies of every glycosylated clone.

**Figure 2:**
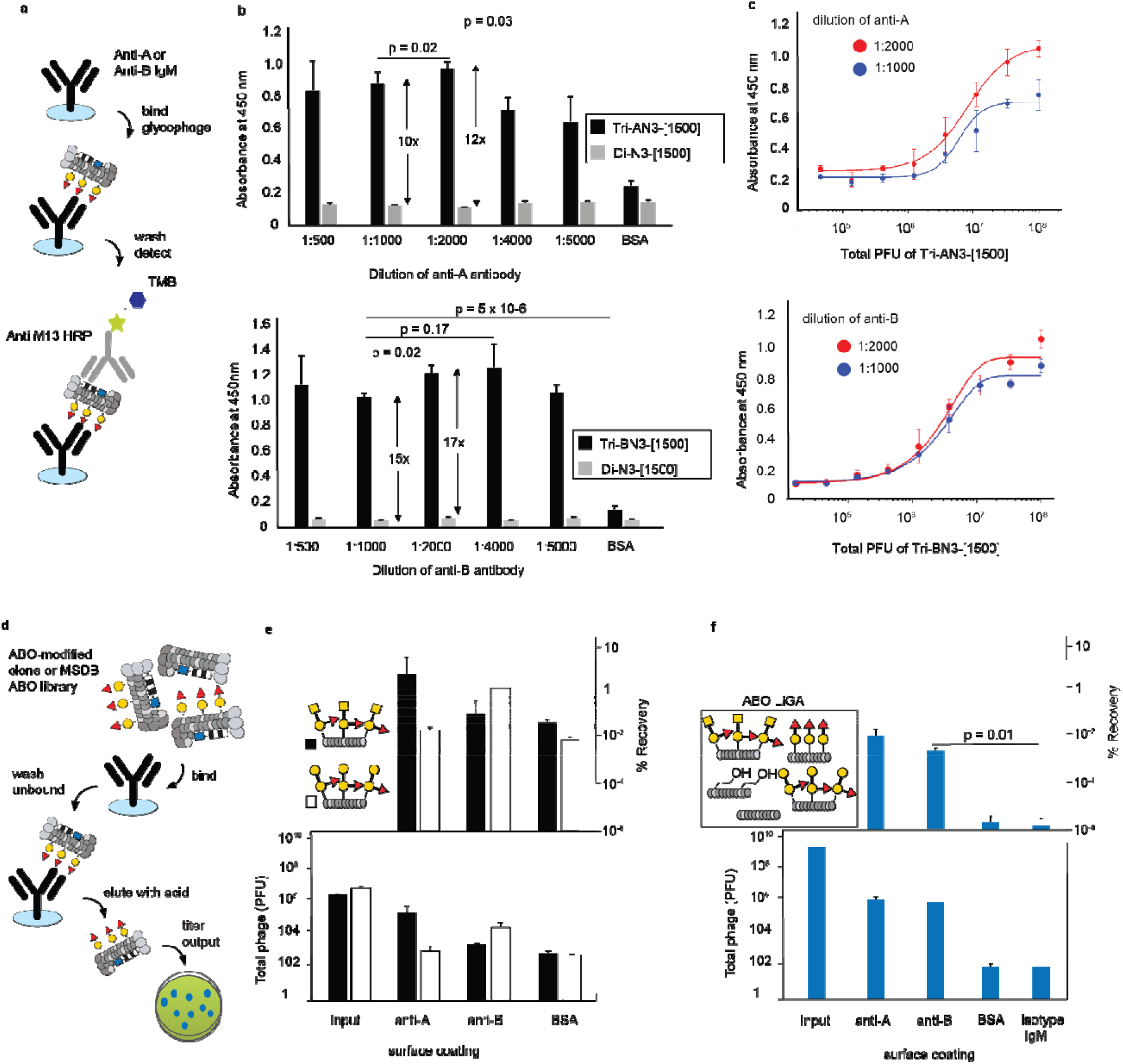
Functional validation of ABO-phage conjugates using ELISA. **a)** Representation of the phage-ELISA assay. **b)** Dilution studies to determine the optimum coating concentration of anti-A (top) and anti-B (bottom) antibodies: Antibodies were coated at five different dilutions and tested with phage glycosylated with either the A or B antigen. Indicated p-values are calculated using two tailed t test. **c)** EC_50_ of ABO phage particles as determined using dose titrations. **d)** Graphical illustration of functional validation of MSDB phage-ABO glycoconjugates using a plaque forming assay. **e)** Recovery of phage clones glycosylated with Tri-AN3 and Tri-BN3 on anti-A, anti-B antibodies and BSA coated wells. **f)** Recovery of MSDB-ABO LiGA library on anti-A, anti-B antibodies, isotype mouse IgM and BSA coated wells.

We further performed a plaque forming unit (PFU) assay to validate the binding of matched and mismatched ABO-glycophages to purified anti-A and anti-B IgM antibodies and controls (**Figure 2d**). The recovery in each experiment was calculated as PFU(output)/PFU(input). We observed that Tri-AN3[1500] exhibited stronger binding to BSA coated wells than Tri-BN3[1500] (**Figure 2e**). In a mismatched binding test, binding of Tri-AN3[1500] to anti-B antibody was higher than binding of Tri-BN3[1500] anti-A antibody (**Figure 2f**). An MSDB ABO-LiGA library that contained five sets of phage conjugates – Tri-AN3[1500], Tri-BN3[1500], Di-N3[1500], azidoethanol and non-conjugated “blank” phages were mixed and tested on anti-A and anti-B antibodies and an isotype mouse IgM antibody as a control. The library recovery on anti-A antibody was 10,000-fold higher than that on the isotype antibody, whereas the recovery on anti-B antibody was 8,000-fold higher when compared to that of the isotype antibody (**Figure 2f**). Combined MALDI, ELISA and PFU assay validated the structural and functional integrity of the A and B trisaccharide antigens on the phage, and their ability to bind IgM anti-A and anti-B. These experiments provided evidence for weak, but detectable, non-specific interactions between the Tri-AN3[1500] construct and BSA.

### One-step vs. two-step PCR for amplification of LiGA library

Amplification of phage samples obtained from LiGA binding assays on anti-A and anti-B antibodies can be done either by a one-step PCR protocol that employs primers with overhangs that contain Illumina adapters (Matochko et al. 2014; Sojitra et al. 2021) or an improved nested two-step PCR protocol (Sojitra et al. 2025) in which the first step is performed by 18–20 nt primers with no overhang (**Figure S5**). The primers used for the first step of the two-step PCR yielded robust linear responses between Cq values in qPCR assays and copy number of phage for 10–100 copies (**Figure S6**). On the other hand, primers used for one-step PCR frequently failed for phage samples that had a titer of <10^4^ copies (**Figure S7a**): Amplification of these samples produced “primer-dimers” with no detectable amplification of the target sequence. In contrast, the two-step PCR method resulted in successful amplification of samples with a copy number of 5,000 (**Figure S7c**).

Numerous publications from our lab successfully employed one-step PCR for NGS analysis (Matochko and Derda 2013; Matochko et al. 2014; Sojitra et al. 2021). We reconfirmed these observations using NGS analysis of a model ABO-LiGA processed by one-step vs. two-step PCR. As input for PCR, the total copy number of the glycophages, components of LiGA, was chosen to be 10^6^ because this number can be successfully amplified by both one-and two-step PCR. We were pleased to observe that comparison of the NGS data produced from one-step <u>and</u> two-step PCR yielded correlation with r^2^ of 0.98 (**Figure S8**). Although one-step PCR works in optimal conditions, its efficiency is likely to deteriorate as the amount of phage decreases significantly below 10^6^ copies. The amount of retained LiGA clones on weakly-binding control targets (*e*.*g*., BSA) can fall to the 10^3^–10^4^ PFU range. Also, even in optimal conditions, LiGA contains a mixture of high-copy and low-copy components. Indeed, we noted a systematic deviation between one-step and two-step PCR for low copy number clones (**Figure S8g**). We reasoned that two-step PCR is likely to exhibit fewer biases towards low-copy number constituents when compared to one-step PCR. Two-step PCR was employed in the remaining assays.

### Binding of ABO_1_-LiGA to anti-A and anti-B antibodies

A model ABO-LiGA (ABO_1_-LiGA) combined Tri-AN3, Tri-BN3 and Di-N3 blood group glycans conjugated at one density, together with azidoethanol and a non-conjugated blank as controls (**Figure S4**). Each component of ABO_1_-LiGA was associated with 10 distinct DNA barcodes (MSDBs). Although all glycophage components of LiGA are mixed in a 1:1:1…:1 ratio, the copy number of the resulting amplicons after Illumina sequencing deviated from this ratio (**Figure S9**). We attributed this recurrent observation to poorly understood biases in NGS, which combine: (i) conversion of phage samples to NGS-compatible dsDNA by PCR, (ii) bridge amplification of dsDNA to form clusters on the Illumina chip; (iii) sequencing by synthesis; and (iv) cluster detection and post-processing. We further tested whether the natural dispersion in copy number influences the outcomes from “iso-molecular” observations. In this respect it should be noted that the 10 SDBs are combined to form the MSDB mixture prior to any glycan modification; as a result, all 10 barcodes are associated with an identical phage composition.

NGS analysis tested binding of ABO_1_-LiGA to antibodies immobilized either by adsorption on polystyrene 96-well plates (plate panning) or on Protein-L magnetic beads (bead panning). In plate panning, bound phages were recovered by elution with 10 mM HCl and in bead panning, boiling the beads released the DNA of the bound phage. Differential Enrichment (DE) analysis of the Next Generation Sequencing (NGS) data confirmed the binding of Tri-AN3 and Tri-BN3 to cognate antibodies in plate panning and attenuated enrichment in bead panning (**Figure S9**). Plate panning yielded a fold enrichment (FC) factor of >5 above the background and a Z’ score of 0.57 for the anti-B antibody and 0.39 for the anti-A antibody, indicating robust signal by HTS metrics. In bead panning, the FC decreased to <2. According to Bioconductor-DE criteria binding to cognate antibodies was statistically significant for all ten clones bearing Tri-AN3 and 8/10 clones bearing Tri-BN3. However, Z’ scores of 3.1 and –1.1 in bead panning experiments suggested poor assay performance (**Figure S9**). We note that, unlike optimized surface loading for plate panning (**Figure 2b**), we did not systematically optimize loading of antibodies on Protein L magnetic beads. We also did not test an alternative support (*e*.*g*., anti-IgM-coated beads) nor tune the wash stringency of beads: both can influence the outcome of a LiGA assay. If necessary, such optimization can be conducted in the future. For the purposes of this manuscript, we performed all remaining experiments using by antibodies adsorbed on plates.

We mixed ABO_1_-LiGA with the previously published LiGA (Sojitra et al. 2021) to create an expanded LiGA (designated here as ABO_2_-LiGA) with 65 different glycophage constructs including the A and B subtype I and II tetrasaccharide antigens. The 65 other glycosylated phage clones did not influence binding of Tri-AN3 and Tri-BN3 to cognate antibodies (**Figure S10**). In an anti-B:ABO_2_-LiGA experiment, enrichment was observed for not only for Tri-BN3[1500] but also for the B type VI (B-tetra-L)[920] and the B type I tetrasaccharide[350] but not for the B type II tetrasaccharide[1000]. In an anti-A:ABO_2_-LiGA experiment, enrichment was observed for Tri-AN3[1500] and Tri-AN3[860], a component of LiGA-65, but not for the A type I tetrasaccharide[350] nor A type VI tetrasaccharide (A tetra L) (**Figure S10**).

Both purified GBPs and GBPs expressed on cell surfaces (Sojitra et al. 2021) require a specific density of glycan on phage for effective recognition. To test the effect of ABO glycan density on their recognition by the antibodies, we produced ABO_3_-LiGA in which glycans were displayed at low (90– 120 copies), medium (500–700 copies) or high (1500–1800 copies) densities confirmed a critical role for the high density of glycans. Neither Tri-BN3[100] nor Tri-BN3[540] exhibited any binding to anti-B IgM, but in the same ABO_3_-LiGA phage decorated with high copy number Tri-BN3[1500] bound strongly to anti-B and differentiate anti-B from anti-A with a Z’ score of 0.5 (**Figure 3**). Tri-AN3[1500] bound strongly to anti-A and differentiate anti-A from anti-B with a Z’ score of 0.82. In the same mixture, the Tri-AN3[540] response was detectable but severely dampened and the binding of Tri-AN3[100] to anti-A IgM was very weak across all 10 SDBs. These observations suggest that serological assays critically depend both on glycan density on phage and on presentation of antibodies in the assay.

**Figure 3:**
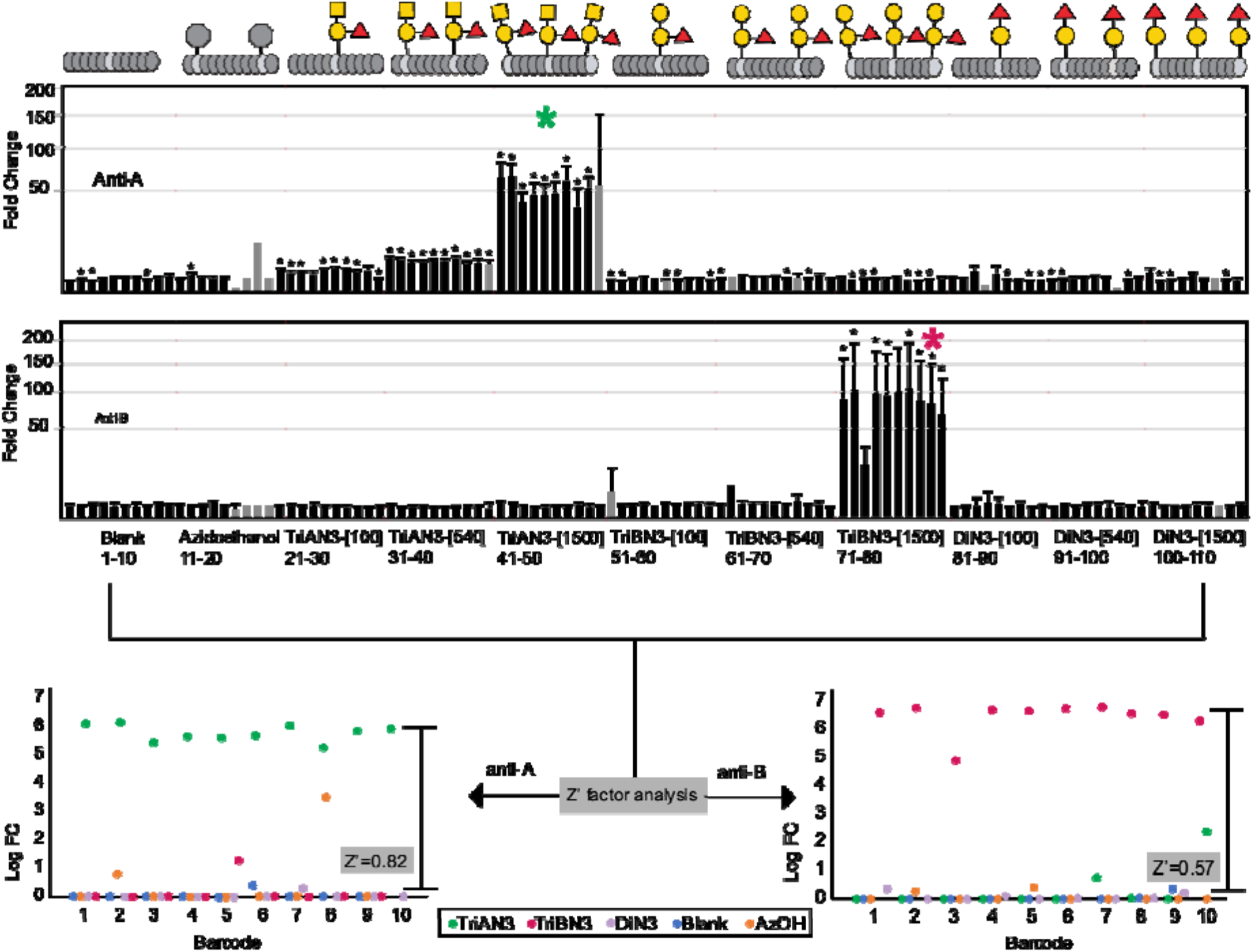
Density scan of MSDB ABO_3_-LiGA anti-A and anti-B antibodies and Z’ score analysis of MSDB ABO_3_-LiGA screening on anti-A anti-B antibodies. Error bars designate variance propagated from the variance of the NGS data for input and output populations. Asterisks indicate significance of enrichment over median population.

Comparison of the data presented in **Figures 3, S9** and **S10** yields the curious observation that a LiGA with more components has higher fidelity. The FC values for anti-A–Tri-AN3 and anti-A:Tri–AN3 interaction increased from FC=5 in ABO_1_-LiGA (**Figure S9**) to FC=20 in ABO_2_-LiGA (**Figure S10**) to FC=50-100 in ABO_3_-LiGA (**Figure 3**). We believe that this increase is a consequence of Bioconductor-DE processing designed for DE-analysis of RNAseq data. Specifically, the algorithm assumes the existence of an endogenous median population of “non-responding” transcripts, which is employed as a baseline for calculating FC values. A LiGA mixture that contains many non-binding clones, in principle, adheres to such an assumption. Bioconductor-DE assumptions may fail if the LiGA composition is limited or if most LiGA components are binders, as it might happen in a serological assay. For such cases, a prospectively determined *bona fide* baseline must exist in all LiGA experiments. In previous publications, we employed unmodified, “blank” (or wild type, wt) phage, or phage in which DBCO had been capped by azidoethanol, as the baseline. The data in **Figure 3** suggest that both wt and azidoethanol (FC=1), in theory, might be useful as an *a priore* baseline in downstream serological tests. To further reconfirm these observations, we compared two array compositions— ABO_3_-LiGA and an expanded ABO_4_-LiGA created by combining ABO_4_-LiGA with a previously published set of glycosylated phages (Sojitra et al. 2021)—in assays that employed IgG and IgM repertoire from whole serum.

### Testing of ABO LiGA on human serum samples

To test the ABO_3_-LiGA and ABO_4_-LiGA in serological assays, we employed serum samples from 31 healthy donors and tested both IgM and IgG antibodies to ABO blood group determinants (Bentall et al. 2021; Jeyakanthan et al. 2016; Saddam M. Muthana and Gildersleeve 2016b). As a benchmark, we used a Luminex-based ABO antibody assay that has been validated with a larger panel of 143 individuals (O: n=68; A: n=48; B: n=17; AB: n=10, Anne Halpin, “Immune Risk Assessment in Pediatric Heart Transplant Patients” <u>PhD Thesis</u>) and 134 individuals (O: n = 67; A n = 51; B: n = 16) (A. M. Halpin et al. 2025). A set of 31 samples have been tested both by Luminex and LiGA. In short, ABO Luminex employs, A- and B-trisaccharides and 12 tetrasaccharide subtypes—six for each A and B, in addition to H-disaccharide and six H-trisaccharide subtypes -—conjugated to individual beads. The glycan-coupled beads were incubated with patient plasma/serum, washed to remove unbound antibodies and probed with phycoerythrin (PE)-labelled IgG- and IgM-specific secondary antibodies. The mean fluorescence intensity (MFI) for each bead was detected using a Luminex 200 system (**Figure 4A**). LiGA similarly tested the glycan reactivity of IgG and the IgM antibodies. Serum samples at 1:100 dilution (n=4 for each sample) were by immobilized in a 96-well plate coated with either anti-human IgM or anti-IgG. ABO_3_-LiGA or ABO_4_-LiGA was added to the plate, washed following a previously optimized wash protocol (**Figure 2-3**), recovered by acid elution, amplified by two-step PCR and analyzed by NGS to determine the FC enrichment of each glycan. From 31 available samples, sample recovery or PCR failed for 8/31 IgG and 9/31 IgM samples (see grey bars in **Figure 4B**); the other samples yielded high-quality NGS data and are summarized in **Figure 4B**. For simplicity, the data in **Figure 4b** is a median value from 10 MSDB and **Figure S11** summarizes the same data prior to averaging responses across MSDB. We noted that although ABO_3_-LiGA and ABO_4_-LIGA differ in complexity, measurement of seven samples with the both ABO_3_-LiGA and ABO_4_-LiGA (**SI Figure S10 and S12A**) demonstrated that binding of AN3 and AB3 and diN3 to the IgM repertoire was reproducible and was not influenced by LiGA composition (**Figure S12-S13**). Based on this observation, we aggregated the data from both observations; calculated the median from the observations where both datasets were available and designated these observations as “ABO_M_-LIGA” in the discussion below.

**Figure 4.**
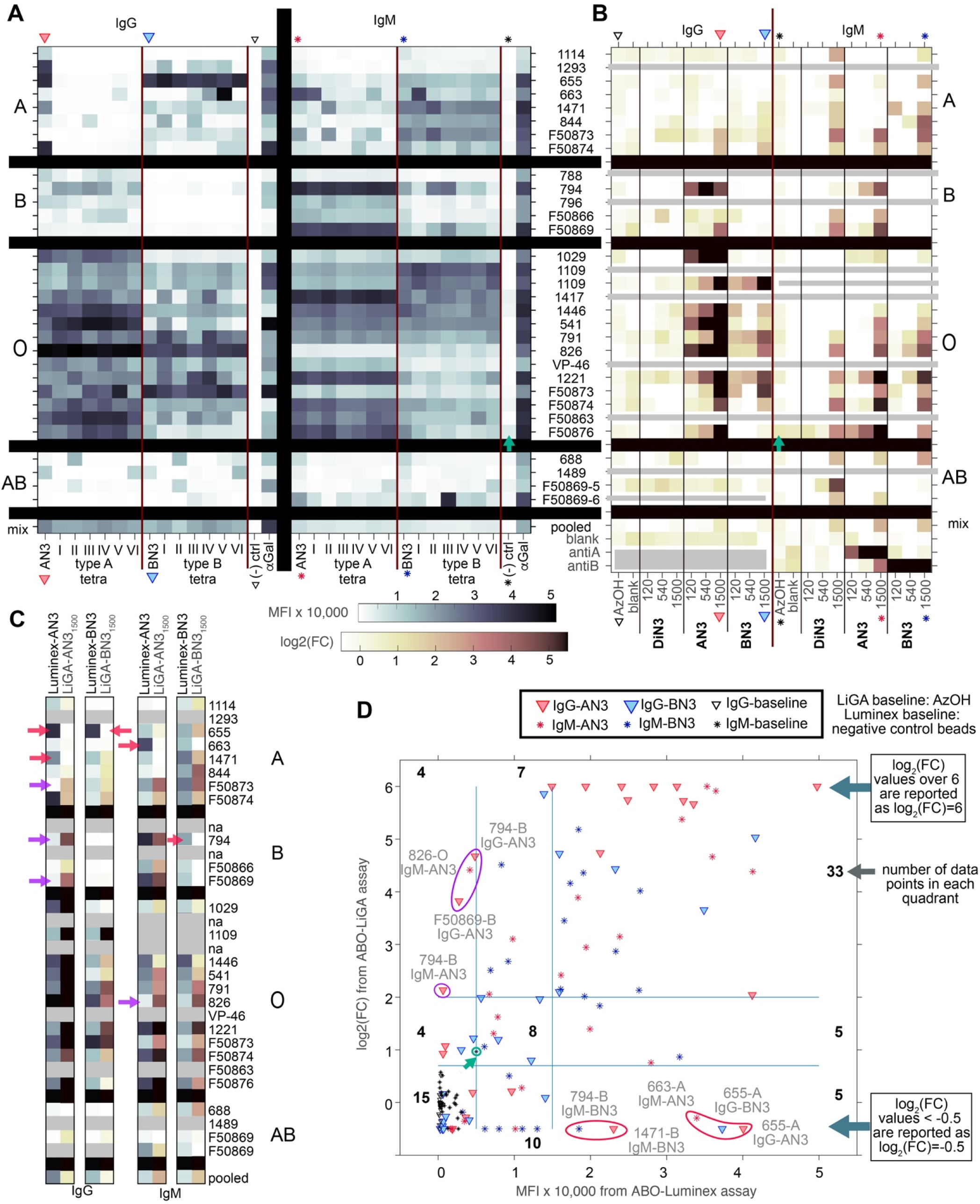
Comparison of the ABO-Luminex and ABO_M_-LiGA for 31 serum samples. **A)** Heatmap with the mean MFI of IgG and IgM binding to Luminex beads bearing A or B type tri- and tetrasacharides. **B)** FC enrichment of ABO_M_-LiGA glycans on IgG and IgM-coated surfaces. ABO_M_-LiGA employs three densities of AN3, BN3, and diN3. LiGA data is a median of measurements from 8–10 DNA-encoded MSDB replicates for each glycan. For unaveraged data and dispersion in the data due to PCR amplification or NGS analysis, see the supporting information. Data acquisition failed for 8/31 IgG and 9/31 IgM samples (grey bars). **c)** Comparison of the AN3 and BN3 data for IgG and IgM repertoires, color intensities are identical to those in panels A or B. In **A–C**, data is clustered by blood type in the rows and by glycans in the columns. D) Scatter plot of MFI from Luminex vs. log2(FC) from LiGA; numbers denote the number of datapoints in each quadrant.

Observations in ABO-Luminex mirror those previously seen in a larger cohort (A. M. Halpin et al. 2025) where over half of the ABO-A individuals had detectable IgG and IgM antibodies to the A-trisaccharide (AN3) but less than 10% of the ABO-A healthy controls exhibited reactivity to A-tetrasaccharides (Anne Halpin, <u>PhD Thesis</u>). In line with previous reports, ABO-Luminex detected high levels of IgG and IgM antibodies reactive to AN3 in 4/8 ABO-A individuals; but all tested ABO-A individuals had comparatively low levels of antibody to A-II, III, and IV subtype glycans (**Figure 4A**). The original ABO_3_-LiGA was built on AN3 and BN3 trisaccharides only; however, the IgG and IgM reactivity to the AN3/BN3 trisaccharides correlates poorly with blood group as determined by the hemagglutination assay (Anne Halpin, <u>PhD Thesis</u>). Still, we reasoned that the comparison of the reactivity of AN3 and BN3 trisaccharides between ABO-Luminex and ABO_M_-LiGA can be used to test whether the performance ABO_M_-LiGA has any substantial deviations from ABO-Luminex. We focused the comparison on Tri-AN3[1500] and Tri-BN3[1500] samples, which exhibited high Z’-scores (**Figure 3**). **Figure 4C** aligns the AN3 and BN3 reactivity to IgG and IgM detected by LiGA and Luminex: the most substantial discrepancies in the reactivity of IgG and IgM towards AN3 and BN3 are highlighted by red and purple arrows.

To further compare both datasets, we plotted the ABO-Luminex *vs*. ABO_M_-LiGA data (**Figure 4D**) and binned the data into background, low, and high signals. In Luminex, we set the background threshold to MFI=3,000 to delineate >95% of non-specific binding of IgG and IgM to negative control beads (vertical thick blue line, **Figure 4D**). In LiGA, a similar log_2_(FC)=0.7 threshold was set to delineate binding of IgM/IgG to phages in which the DBCO reactive group was capped by azidoethanol (see horizontal thick blue line in **Figure 4D**). The purpose of the upper threshold in Luminex and LiGA was to separate strong and weakly reactive samples. We set the upper threshold in Luminex to 15,000 MFI based on previous report of median value of IgG and IgM reactive populations (A. M. Halpin et al. 2025). Setting a threshold of log_2_(FC)>2 in LiGA allowed to classify 15+7+35 = 57 responses into the same category (background, weak or strong binding). Only 2+5 = 7 responses were a complete mismatch. Five samples exhibited only “background” binding in ABO_M_-LiGA but they exhibited strong reactivity in ABO-Luminex (bottom right quadrant in **Figure 4D**). Conversely, two ABO-B samples had only background IgG binding to AN3 in ABO-Luminex but strong binding in ABO_M_-LIGA (upper left quadrant in **Figure 4D**). Adjustment of the MFI=3,000 baseline may exclude or include borderline samples, which are annotated in **Figure 4D**. From 14+7+10=31 weak ABO-Luminex signals, 14 samples had only “background” reactivity in ABO-LiGA. Conversely, ABO-Luminex did not detect 2/12 samples that were weakly reactive in ABO_M_-LiGA. The trends overall suggest that the current version of LiGA fails to detect nearly 45% of samples with weak reactivity in ABO-Luminex; but it detects 38/43=88% of samples with strong reactivity in ABO-Luminex.

A puzzling observation in ABO_M_-LiGA was that IgM exhibited a consistently higher binding to AzOH phage when compared to wt (blank) phage; whereas, IgG binding to AzOH phage was indistinguishable from IgG binding to wt (blank) phage (**Figure 4b**). In previous reports, binding of wt and AzOH to purified antibodies (**Figure 3**), lectins, or GBPs on cells (Sojitra et al. 2021) served as a convenient baseline. In serological experiments, if AzOH are employed as a “baseline” then the binding of AN3-DBCO or BN3-DBCO to IgM is often significantly below such a “baseline”. It appears that the glycans shield the DBCO linker from non-specific binding to the serum antibody repertoire. The binding to DBCO capped by AzOH is modest in general but it might explain rare instances when AN3 or BN3 glycans in ABO_3_-LiGA detected a signal not detectable by ABO-Luminex. Such binding might simply be an artefact of the DBCO-based immobilization strategy used to generate ABO_3_-LiGA. Nonspecific binding of the DBCO linker might also explain the erratic and often strong binding of disaccharide DiN3, which, due to its smaller size compared to AN3 and BN3, may be less effective in shielding serum antibodies from the linker. This DiN3-DBCO conjugate is frequently enriched in other LiGA experiments.

ABO_4_-LiGA contained a limited collection of A and B tetrasaccharides, type I, II and VI, albeit each glycan was displayed at one density and encoded by one DNA barcode. A limited set of glycans, barcodes, and a small number of tested serum samples (**SI Figure S10 and S12A**) prevented in depth comparisons of tetrasaccharide response between ABO_4_-LiGA and ABO-Luminex. Future experiments with a complete set of tetrasaccharides of predetermined density will have to be conducted to expand these observations. In conclusion, the ABO-LiGA assay confirmed observations from the Luminex ABO assay, identified shortcomings when compared to Luminex ABO and pointed to a critical need for two upgrades in the LiGA technology: (1) replacement of AN3/BN3 by A/B tetrasaccharides; (2) comparison of the DBCO linker and another linker (*e*.*g*., amide linker employed in ABO Luminex assay).

## Discussion

This paper demonstrates that LiGA can detect IgG and IgM ABO blood group antibodies in a complex mixture like human serum. Performance of ABO-LiGA based on trisacharide determinants is similar to the performance of the same determinants in the ABO-Luminex bead assay; however, it does not match the robustness of latter better-established assay. In particular, failure of the PCR for 20–30% of the tested serum samples suggests that there is not a single set of assay conditions to measure the response of all serum samples. The dilution, immobilization or washing strategy needs to be tailored for different serum samples. A relatively high false positive rate—antibody reactions detected in ABO-Luminex but not ABO-LiGA—again points to the need to optimized conditions for sample handling in DNA-encoded analysis of interaction of carbohydrates with serum antibodies. A cursory analysis of the effect of adding 65 other phage-linked glycans to the mixture demonstrates that the increased complexity of the LiGA does not influence the outcome. Testing of more complex LiGA mixtures with multi-barcoding and multi-density constructs is a next logical step. Future experiments will also involve an expanded ABH panel with 18 blood group subtypes that mimics previously reported ABH-arrays (Jeyakanthan et al. 2016) and the ABO-Luminex assay (A. M. Halpin et al. 2025).

Errors in PCR or Illumina NGS might be postulated to be one of the reasons for divergence between the LiGA and Luminex assays. However, a predetermined Hamming distance between all barcodes allows their identification, even if the PCR or sequencing procedure introduces random errors. Discussion of sequence-based strategies for error correction can be found elsewhere (Buschmann and Bystrykh 2013; Bystrykh 2012; Hamming 1950; Levenshtein 1966). Difference in the intensity of signals in the two assays might also originate from non-uniform PCR amplification of DNA strands prior to NGS (Heckel et al. 2019). Preferential amplification of some DNA barcodes over others using PCR distorts the distribution of copy number (Pan et al. 2014; Ruijter et al. 2009; Warnecke et al. 1997). The bias is particularly severe for sequences with low copy numbers in the initial mixture due to PCR stochastic bias (Chen et al. 2020). This problem is exacerbated with decreases in copy number of templates (SI Fig. S5). We believe that similar problems are mirrored in this manuscript: A mixture of 10 distinct DNA barcodes (MSDBs) encoding the same glycan allowed us to visualize distinct outliers, which likely originate from similar PCR biases. We envision that discarding outlier(s) SDBs prior to averaging the MSDB response will be a productive strategy for dealing with PCR bias. In addition, the assay might not require as many as 10 barcodes and a smaller number (e.g., 4–7) in which outliers are discarded, is plausible.

Prior to NGS analysis, we employed phage ELISA (Barbas et al. 2004) and PFU assays to evaluate the functional integrity of individual glycosylated phage constructs, to optimize the concentration of purified anti-A and anti-B antibody to be coated on the plate and to fine tune the concentration of the ABO-LiGA library to be used for binding experiments. Despite its age (100 years old) (Anderson et al. 2011) we would like to highlight the value of the PFU assay; its outstanding dynamic range still makes it best-in-class for optimizing phage–receptor interactions. Unlike ELISA or PCR, the PFU assay reliably detects one molecule (particle) of phage in an assay mixture. In a typical assay with a 100 μL volume, this corresponds to 10^4^ molecules/L or a 1.6 attomolar concentration. Such sensitivity cannot be matched by PCR, which suffers from amplification bias at a low dilution of the DNA template, or ELISA, which typically requires many copies of the enzyme for reliable a signal-to-noise ratio to be obtained.

We observed several indications that the DBCO linker may cause non-specific interactions and skew the results of the assay. Problems of this type were never detected in prior investigations, which employed purified lectins. Here, we also showed that DBCO blocked with azidoethanol shows no detectable problems in assays with purified antibodies. This observation reassured us that, using purified proteins, responses in LiGA are not influenced by the linker. However, DBCO-related problems appear to be present in complex mixtures (*e*.*g*., IgM repertoire shows stronger binding to DBCO-azidoethanol when compared to unmodified phage) or in conjugates with small glycans. The diN3 disaccharide exhibits erratic reactivity across all serum samples (**Figure 4B**) but shows no binding to purified antibodies (**Figure 3**). Increasing the glycan size and replacement of DBCO is a critical next step for future investigations. Replacing of the A/B trisaccharide antigens with the A/B tetrasaccharide may also yield improvements in assay performance through enhancing the specificity of binding. Experiments are ongoing to conjugate the expanded set of ABH tetrasaccharides to MSDBs with several different linkers.

## Supplementary Information

Details of the experimental procedures, data processing, supplementary Figures **S1**-**S13**, list of all next generation sequencing files using in this study and instructions for access (**Table S2-S4**)

## Acknowledgements

We thank Jeff Gildersleeve at National Cancer Institute for generous donation of the serum samples for this manuscript and manuscript-related discussions. This research was supported by grants from CIHR (#180445 to R.D.), GlycoNet (CR-29 to R.D.), Alberta Innovates Strategic Research Projects (to R.D.), Alberta Innovates Graduate Student Scholarship (to G.M.L.).

