## Supporting Information for "Evaluation of Multiplexed Liquid Glycan Array (LiGA) for Serological Detection of Glycan-binding Antibodies"

#### Abbreviations

| | |
|---|---|
| ELISA | Enzyme Linked Immunosorbent Assay |
| IgG | Immunoglobulin G |
| IgM | Immunoglobulin M |
| BSA | Bovine Serum Albumin |
| PCR | Polymerase Chain Reaction |
| qPCR | Quantitative Polymerase Chain Reaction |
| NGS | Next Generation Sequencing |
| *E. coli* | *Escherichia Coli* |
| LiGA | Liquid Glycan Array |
| SDB | Silent Double Barcode |
| MSDB | Multiplexed Silent Double Barcode |
| PFU | Plaque Forming Units |
| SPAAC | Strain Promoted Azide–Alkyne Cycloaddition |
| DBCO | Dibenzocyclooctyne |
| MALDI-TOF | Matrix Assisted Laser Desorption Ionization – Time of Flight |
| Tri-AN3 | A Trisaccharide Antigen |
| Tri-BN3 | B Trisaccharide Antigen |
| Di-N3 | H Disaccharide Antigen |
| FC | Fold Change |
| MFI | Mean Fluorescence Intensity |

### 1. Biochemical Methods

**1.1.** Construction of silent distal barcode (SDB) M13 phage library

This protocol is adapted from the previous publication(Tjhung et al. 2016). A library of degenerate codons, termed, silent double barcodes (SDB) was created in the phage genome at a position proximal to the gIII cloning site starting from a M13KE vector containing the stuffer sequence CAG TTT ACG TAG CTG CAT CAG GGT GGA GGT corresponding to the peptide QFT*LHQ, with * representing an amber stop codon. The insert fragment was PCR amplified using the primers **P1** and **P2** and the vector fragment was PCR amplified using primers **P3** and **P4**:

**Name** Sequence (5’->3’):

**P1**GAGATTTTCAACGTGAAAAACT**N**CT**N**TT**Y**GC**N**AT**H**CC**N**CT**N**GTGGTACCTTTCTATTCTCA

**P2** TTAAGACTCCTTATTACGCAGTA

**P3** TTGCTAACATACTGCGTAATAAG

**P4** TTTTTCACGTTGAAAATCTC

**P5**GTGGTACCTTTCTATTCTCACTCGAG**Y**GT**N**GA**R**AA**R**AA**Y**GA**Y**CA**R**AA**R**AC**N**TA**Y**CA**Y**GC**N**GG**N**GG**N**GG**N**T-CGGCCGAAACTGTTGAAAG

**P6** CGAGTGAGAATAGAAAGGTAC

PCR was performed using 50 ng phage dsDNA with 1 mM dNTPs, 0.5 µM primers, 0.5 µL Phusion High Fidelity DNA polymerase in 1x PCR buffer (NEB #B0518S) in a total volume of 50 µL. The temperature cycling protocol was performed as follows:

a) 98 °C 3 min,

b) 98 °C 30 s,

c) 60 °C 30 s,

d) 72 °C 4 min,

e) repeat b) - d) for 35 cycles,

f) 72 °C 10 min,

g) 4 °C hold.

PCR amplified fragments were treated with restriction enzyme DPN 1 (NEB #R0176S) and then gel purified. NEBuilder Hifi DNA assembly was then carried out following the manufacturer protocols by mixing 100 ng of vector, 4 ng insert, 10 µL of NEBuilder Hifi DNA assembly master mix (NEB #E2621S), and deionized H2O up to a total volume of 20 µL. The resulting ligated DNA was transformed into *E.coli* K12 ER2738 and propagated overnight at 37 °C. The overnight culture was then centrifuged to separate bacteriophage from host cells. Deep sequencing of the resulting SB1 QFT*LHQ cloning vector with 6,144 theoretical sequence combinations in the leader region is available at: https://48hd.cloud/file/20161105-68OOooIC-NB. The cloning of a degenerate codon library of the SVEKNDQKTYHAGGG peptide to produce SB2 region was conducted as follows. Primers **P5** and **P2** were annealed to and amplified by PCR following the

35-cycle protocol described above to produce a dsDNA insert. The vector SB1 QFT*LHQ was PCR amplified using primers **P4** and **P6**. PCR fragment were processed using NEBuilder Hifi DNA assembly kit (NEB #E2621S) as described above. The resulting ligated DNA was transformed into electrocompetent cells *E.coli* SS320 (Lucigen). The resulting overnight culture was centrifuged to remove host cells and incubated with 5% PEG-8000, 0.5 M NaCl for 8 h at 4 °C, followed by 15 min centrifugation at 13000 g to concentrate released phage. PEG precipitated phage were re-suspended in PBS-Glycerol 50% and stored at -20 °C. Sequence of the vector containing M13-SDB-SVEKY library is available on GeneBank (#MN865131). Deep sequencing of the resulting library is available at: https://48hd.cloud/file/20161215-67OOooOO-NB

**1.2.** SDB clone isolation and amplification

Isolation of SDB clones was performed as described in previous publication(Sojitra et al. 2021) from M13-SDB-SVEKY library. A 10 µL aliquot of phage was diluted and plated at a density of 100 plaques per plate. Single colonies were manually picked, and individually transferred into a clean 1.7 mL plastic tube containing 0.5 mL of PBS-Glycerol 50% and incubated at room temperature for 30 min. The tubes were then placed in 55 °C heating block for 10 min to inactivate any remaining bacterial cells. After the incubation, 20 µL sample of each colony suspension was amplified for 4.5 h in 5 mL of LB supplemented with a 0.5 mL of log phase *E. coli* K12 ER2738. After amplification the phage clones were collected from the culture supernatant by centrifugation at 4500 g for 10 min. Next, bacterial cell pellet and culture supernatant were processed separately. The supernatant was incubated with 5% PEG-8000, 0.5 M NaCl for 8 h at 4 °C, followed by 15 min centrifugation at 13000 g to precipitate the viral particles. The phages were re-suspended into 1 mL PBS-Glycerol 50%, titered, and stored at -20 °C until further use. The bacterial cell pellet was processed for phage-DNA extraction using GeneJET Plasmid Miniprep kit (Thermo Fisher, #K0502). For SDB identification, a sample of 400 ng of the phage DNA was submitted for Sanger sequencing. We selected the clones that contained three or more base pair substitutions from one another (i.e., Hamming distance (H)≥3). H≥3 permits correction of any point mutations that may have arisen during the analysis by deep sequencing.

**1.3.** Construction of Multiplex Silent Distal Barcodes (MSDBs)

There are two ways to construct MSDBs (Fig. S2). In the first method, an MSDB was prepared by mixing 10 phage clones each with a unique SDB. Individual phage clones amplified in 25 mL flasks were characterized by deep sequencing and mixed by matching the titer of the phage stock. The MSDB was then used to conjugate blood group antigens. Each MSDB was assigned a two-letter identifier (e.g., ZA) and a "dictionary" (e.g., ZA.xlsx, Fig. S3), a table that describes the correspondence between the DNA barcodes and the glycan conjugated to the respective MSDB. In the second method, 10 individual clones were amplified separately in 3 mL volume and mixed together before purification of phage by PEG precipitation. The initial characterization of MSDBs described in this publication were constructed using the first method, however, since the second method utilizes lesser volume of reagents (amplification in 3 mL tubes as opposed to 25 mL flasks in the first method), the majority of the MSDBs were constructed using the second method.

**1.4.** Ligation of ABO glycans to M13 phage

A solution of SDB phage clone ($10^{12}$-$10^{13}$ PFU/mL in PBS) was combined with DCBO-NHS (20 mM in DMF) to afford a 0.2-2.0 mM concentration of DCBO-NHS in reaction mixture, which typically yields 5-50% of pVIII modification after 45 min of incubation. After conjugation of DBCO-NHS, each clone was individually purified on a Zeba™ column following the manufacturer instructions. Solutions of azido-glycans (Han et al. 2020) (10 mM stock in Nuclease Free $H_2O$) were added to the filtrates to afford a 2 mM concentration of glycan-azide and the solutions were further incubated overnight at 4 °C. All chemical reactions were verified and quantified by MALDI-TOF as described on the previous sections. If reactions were incomplete and residual pVIII-DBCO peak was detected in MALDI, we added additional amount of azido glycan and extended the incubation. If reactions were completed, the conjugates were purified by Zeba ™ column and stored at 4 °C or supplemented by glycerol and stored as 50% glycerol stock at -20 °C. LiGA mixture was prepared by combining these stock solutions.

**1.5.** Characterization of phage-ABO glycoconjugates by MALDI-TOF MS

The sinapinic acid matrix was formed by deposition of two layers. Layer 1 was prepared as 10 mg/mL solution of sinapinic acid (Sigma, #D7927) in acetone-methanol (4:1). Layer 2 was prepared as 10 mg/mL solution of sinapinic acid in acetonitrile:water (1:1) with 0.1% TFA. In a typical sample preparation, 2 µL of phage solution in PBS was combined with 4 µL of layer 2, then a 1:1 mixture of (layer 1): (layer 2+phage) was deposited in that order onto the MALDI inlet plate ensuring that layer 1 is completely dry before adding layer 2+phage. The spots were washed with 10 µL of water with 0.1% TFA to remove salt ions from the PBS. To estimate the ratio of modified to unmodified pVIII, we fit and plotted the data using MatLab.

**1.6.** Methods that measure binding of LiGA components

**1.6.1.** Measuring binding to dilutions of anti-A and anti-B antibodies with ELISA

Murine monoclonal anti-A and anti-B IgM antibodies (produced by Immucor, Inc. and generously donated by Dr. Lori West at the University of Alberta) were each dissolved in PBS to yield the dilutions – 1:500, 1:1000, 1:2000, 1:4000 and 1:5000. 50 µL of the dilutions was added to each well of 96 well plate (Corning®, #CLS3369). The plate was covered with sealing tape (Thermo Scientific, #15036) and incubated overnight at 4 °C. The following day, the wells were washed 3 times by adding washing buffer (200 µL, 0.1% Tween-20 in PBS) in the wells and discarding the solution by inverting the plate on top of a paper towel. Thereafter, blocking solution (100 µL, 20 µg/uL BSA in PBS) was added to the wells and incubated for 1 h at rt. The solution was discarded by inverting the plate, the wells were then washed three times with washing buffer. The wells coated with anti-A and anti-B antibody respectively and incubated for 1.5 h at rt. The solution was discarded by inverting the plate, the wells were washed three times with washing buffer (200 µL 0.1% Tween-20 in PBS) and probed with anti-M13-HRP conjugate. 100 µL of TMB ELISA substrate (Thermo Scientific, #34028) was added to each well and incubated for 5-10 min until a deep blue color developed. The solutions were then quenched with 100 µL 1 M phosphoric acid and absorbance at 450 nm was recorded.

**1.6.2.** Binding of glycosylated clones to anti-A and anti-B measured by ELISA

Blood group reagents Anti-A and Anti-B murine monoclonal antibody (Obtained from Immucor, Inc. and generously donated to me by Dr. Lori West at the University of Alberta) was dissolved in PBS at final dilution of 1:1000 and 1:2000. 50 µL of the solution was added to each well of 96 well plate (Corning®, #CLS3369). The plate was covered with sealing tape (Thermo Scientific, #15036) and incubated overnight at 4 °C. The following day, the wells were washed 3 times by adding washing buffer (200 µL, 0.1% Tween-20 in PBS) in the wells and discarding the solution by inverting the plate on top of a paper towel. Thereafter, blocking solution (100 µL, 20 µg/uL BSA in PBS) was added to the wells and incubated for 1 hour at rt. The solution was discarded by inverting the plate, the wells were then washed three times with washing buffer. Thereafter, solutions of LiGA clones (50 µL in PBS) were added to the wells and incubated for 1 hour at rt. The solution was discarded by inverting the plate, the wells were washed three times washing buffer (200 µL 0.1% Tween-20 in PBS) and probed with anti-M13-HRP conjugate.

**1.6.3.** Dose titration study of ABO glycosylated clones measured by ELISA

Anti-A and anti-B antibodies were each dissolved in PBS to yield final dilutions of 1:1000 and 1:2000. Plates were prepared as described in previous section **1.4.9.** A antigen and B antigen glycosylated phage clones of concentrations - $10^4$ , $10^5$, $10^6$, $10^7$, $10^8$ PFU were added to wells coated with anti-A and anti-B antibody respectively and incubated for 1.5 h at RT. The solution was discarded by inverting the plate, the wells were washed three times with washing buffer (200 µL 0.1% Tween-20 in PBS) and probed with anti-M13-HRP conjugate. 100 µL of TMB ELISA substrate (Thermo Scientific, #34028) was added to each well and incubated for 5-10 min until a deep blue color developed. The solutions were then quenched with 100 µL 1 M phosphoric acid and absorbance at 450 nm was recorded using BioTek Cytation 5 Plate reader.

**1.6.4.** Binding assay of mLiGA on 96 well plates

A 1:1000 dilution of murine monoclonal IgM anti-A and anti-B antibodies (produced by Immucor Inc. and donated by Lori West at the University of Alberta) was added to a 96-well plate (Corning, CLS3369), which was sealed with tape (Thermo Scientific, 15036) and incubated overnight at 4 °C. The following day, wells were washed three times with washing buffer, incubated with blocking solution (1 h at room temperature) and washed three times with washing buffer. Solutions of m-LiGA was added to the wells (50 µl, $10^{10}$ PFU/mL in PBS) and incubated for 1.5 h at room temperature. Wells were washed three times with washing buffer. To elute bound phage, 50 µl HCl (pH 2.0) was added to the wells and incubated for 9 min at room temperature, and the content of each well was transferred to a microcentrifuge tube containing 25 µl 5× Phusion HF buffer (NEB, M0530S). The neutralized solution was used for titering and as template for the PCR reaction.

**1.6.5.** Binding assay of mLiGA on Protein L beads

A 30 µL suspension of Protein L magnetic beads (Thermofisher, 88850) was rinsed with wash buffer (1 mL). 1 µL of anti-A and anti-B antibody was added to the bead suspension in PBS buffer (1 mL). The mixture was incubated overnight at 4 °C with mixing. The following day, the bead suspension and other reagents were added to a 96-well Deepwell plate (Thermo Fisher, 95040450), and the KingFisher Duo Prime Purification System was used to perform the experiment with the following steps: (1) wash buffer (1 mL), (2) blocking buffer (1 mL), (3) binding with m-LiGA (1 mL), (4) wash buffer (1 mL), (5) wash buffer (1 mL) and (6) PBS buffer. Beads were transferred to microcentrifuge tubes and centrifuged (500g for 1 min). Pelleted beads were resuspended in nuclease-free water (30 µl); 2 µl of the suspension was used in the PFU assay. The remaining solution was incubated at 90 °C for 15 min and centrifuged at 21,000g for 10 min, and 25 µl of the supernatant was used as template for the PCR reaction.

**1.7.** PCR Protocol Section

**1.7.1.** One-step PCR Protocol and Illumina Sequencing

This protocol is adapted from the publication(Sojitra et al. 2021). 25 mL of DNA template solution after bead panning procedure in Nuclease free water and 15 mL of eluted phage after plate panning procedure was amplified in total volume of 50 mL with 1x Phusion® buffer, 50 mM of each dNTPs, 500 mM $MgCl_2$, 1 mM forward barcoded primer, 1 mM reverse barcoded primer and one unit of Phusion® High-Fidelity DNA Polymerase (NEB, #M0530S). In amplification of the naïve libraries, volume of template (phage solution) was 2 µL. Cycling was performed using the following thermocycler settings: a) 98 °C 3 min, b) 98 °C 10 s, c) 50 °C 20 s, d) 72 °C 30 s, e) repeat b)-d) for 10 cycles, f) 98 °C 10 s, g) 72 °C 30s, h) repeat f)-g) for 20 cycles, h) 72 °C 5 min, i) 4 °C hold. The PCR products were quantified by 2% (w/v) agarose gel in Tris-Borate-EDTA buffer at 100 volts for ~35 min using a low molecular weight DNA ladder as standard (NEB, #N3233S). PCR products that contain different indexing barcodes were pooled allowing 10 ng of each product in the mixture. The mixture was purified by eGel, quantified by Qubit (Thermo Fisher) and sequenced using the Illumina NextSeq paired-end 500/550 High Output Kit v2.5 (2x75 Cycles). Data was automatically uploaded to BaseSpace™ Sequence Hub. Processing of the data is described in section "*Processing of Illumina data*".

**1.7.2.** Two-step semi-nested PCR

15 mL of eluted phage after plate panning procedure was used as qPCR template and amplified in total volume of 50 mL with 1x Phusion® buffer, 50 mM of each dNTPs, 500 mM $MgCl_2$, 1 mM forward qPCR primer, 1 mM reverse qPCR primer and one unit of Phusion® High-Fidelity DNA Polymerase (NEB, #M0530S). In amplification of naïve libraries, volume of template (phage solution) was 2 µL. Bio-Rad C1000 qPCR machine was used for DNA amplification. Cycling was performed using the following thermocycler settings: a) 95 °C 3 min, b) 95 °C 10 s, c) 58 °C 30 s, d) 72 °C 20 s (with picture), e) 80 °C 20 s (with picture) f) repeat b)-e) for 34 cycles, g) Melt curve 65 °C – 95 °C (with picture) h) end. qPCR data was analyzed using Bio-Rad CFX Manager software. For the second-step semi-nested PCR, 1-5 µL of the qPCR product was used as DNA template and amplified in total volume of 50 mL with 1x Phusion® buffer, 50 mM of each dNTPs, 500 mM $MgCl_2$, 1 mM forward qPCR primer, 1 mM reverse qPCR primer and one unit of Phusion® High-Fidelity DNA Polymerase (NEB, #M0530S). Amplification was performed using the following thermocycler settings: a) 95 °C 3 min, b) 95 °C 10 s, c) 58 °C 30 s, d) 72 °C 20 s e) repeat b-d for 9 cycles f) 72 °C 5 min g) hold 4 °C. The PCR products were quantified by 2% (w/v) agarose gel in Tris-Borate-EDTA buffer at 100 volts for ~35 min using a low molecular weight DNA ladder as standard (NEB, #N3233S).

**1.8.** Microarray fabrication and binding assay

**1.8.1.** Luminex array

A and B antigens of subtypes I-VI (Meloncelli and Lowary 2010; Meloncelli et al. 2011), as well as the A trisaccharide, B trisaccharide and H(O) disaccharide(Han et al. 2020)antigens were conjugated to Luminex xMAP® beads(Halpin et al. 2025) and quantified using monoclonal ABO antibodies. BSA and αGal antigen were coupled as negative and positive controls, respectively. The detailed protocol on coupling antigens to Luminex beads can be found in the Luminex xMAP cookbookLuminex xMAP Cookbook https://cdn2.hubspot.net/hub/128032/file-213083097-pdf/Luminex-xMap_Cookbook.pdf. IgG and IgM antibodies in sera with specificities for ABO glycans were probed using PE-conjugated Goat Anti-Human IgG and PE-conjugated Goat Human IgM respectively. IgG and IgM signals were recorded as mean fluorescence intensities (MFI).

##### **1.8.2.** MSDB ABO LiGA array

The MSDBs were mixed in a 1:1:1…..:1 ratio based on the phage titer obtained by PFU assay. 1500, 540 or 120 copies of Tri-AN3, Tri-BN3 or Di-N3 was conjugated to each MSDB. Azidoethanol conjugated MSDB and non-conjugated blank MSDB were used as negative controls. LiGA-65 consisting of phage clones with 65 glycans was constructed in the previous publication(Sojitra et al. 2021). MSDB ABO LiGA was mixed with LiGA-65 uniformly based on titer. AffiniPure Goat Anti-Human, Fcγ fragment specific IgG (Cat# 109-005-008) and AffiniPure Goat Anti-Human, Fc5μ fragment specific IgM (Cat # 109-005-043) were purchased from Jackson Immunoresearch Laboratories. IgG and IgM antibodies were diluted to 20 μg/ml in 1X PBS and 50 μl of the dilution solution was added to a 96-well plate (Corning, CLS3369), which was sealed with tape (Thermo Scientific, 15036) and incubated in at room temperature for 2 h. After 2 h, wells were washed two times with washing buffer, incubated with blocking solution (1% BSA, 1 h at RT) and washed three times with washing buffer. Serum samples were diluted (1:100) in 1X PBS and 50 μl was added to wells and incubated overnight at 4 °C. Next day, wells were washed two times with wash buffer. Solutions of MSDB ABO LiGA or MSDB ABO LiGA with LiGA-65 (50 μl $10^8$ PFU/ml in PBS) were added to the wells and incubated for 1.5 h at RT. Wells were washed three times with washing buffer. To elute bound phage, 50 μl HCl (pH 2.0) was added to the wells and incubated for 9 min at room temperature, and the content of each well was transferred to a microcentrifuge tube containing 25 μl 5× Phusion HF buffer (NEB, M0530S). The neutralized solution was used for titering and as template for PCR reaction.

#### 2. Data Processing Methods

##### **2.1.** Processing of Illumina data

This protocol is adapted from the publication(Sojitra et al. 2021). The Gzip compressed FASTQ files were downloaded from BaseSpace™ Sequence Hub. The files were converted into tables of DNA sequences and their counts per experiment, essentially as previously described(Matochko et al. 2014). Briefly, FASTQ files were parsed into separate files based on unique multiplexing barcodes within the reads. Reads that did not contain an identifiable multiplex barcode were discarded. Reads that contained a Phred=0 quality score in any position were also discarded. Additional quality control was based on: (i) mapping the forward and reverse primer regions allowing no more than one base substitution each, (ii) alignment of the forward and reverse read-end overlap, allowing no mismatches between forward and reverse read in the overlap region. Reads that did not match criteria (i) and (ii) were discarded. The two ends of read-pairs that pass the filtering criteria were joined and trimmed to the DNA sequences located between the priming regions; the reads were organized in a tab-delimited text file containing the unique DNA sequences and their copy numbers. Technical replicates were often combined in the same file. Using an SDB-lookup table, we mapped DNA sequences to SDB and translated them to glycans using a LiGA-specific lookup table ("LiGA dictionary"). Any reads with one substitution away from the expected SDB sequence were assigned to the parent SDB sequence. Abbreviated names of glycans are based on those used on CFGwebsite:(http://www.functionalglycomics.org/static/consortium/resources/resourcecored2.sh

tml). The files with DNA reads, raw counts, and mapped glycans were uploaded to http://ligacloud.ca/ server. Each experiment has a unique alphanumeric name (e.g., **20180711-87YOrdRB-JM**) and unique static URL: for example, http://ligacloud.ca/searchLibInfo?f=0&b=0&d=20180711-87YOrdRB-JM. Comparisons and testing differences for significance in the LiGA data were performed essentially as described in our previous publication(Matochko et al. 2014) using DE implemented in edgeR. We used Bioconductor EdgeR DE analysis with modeling of the observed counts using a negative binomial model, Benjamini–Hochberg (BH) adjustment to control the FDR at $\alpha = 0.05$ and normalization of data across multiple samples using the TMM normalization.

#### 3. Origin of Serum

All serum samples were previously collected by Lori West group at the University of Alberta and the Jeff Gildersleeve group at the NIH. Supplementary Table S1 summarizes information regarding the serum samples. Samples from the West group were collected from healthy volunteers at the University of Alberta hospital. Healthy controls gave consent in the Cardiac Transplantation in Infancy, Childhood, and Beyond study, University of Alberta Human Research Ethics Board, Study ID Pro00001408.

Healthy donor sera from the Gildersleeve group were purchased from Valley Biomedical Products and Services (Winchester, VA). The reference pooled human sera were purchased from SeraCare Life Sciences (Milford, MA). Samples were stored in -20°C prior to use.

| Sample ID | Blood type | Age | Sex | Group |
|---|---|---|---|---|
| F50873-02 | O | 49 | M | Gildersleeve |
| F50869-05 | AB | 39 | F | Gildersleeve |
| F50874-05 | O | 20 | F | Gildersleeve |
| F50876-06 | O | 46 | F | Gildersleeve |
| F50873-03 | A | 42 | M | Gildersleeve |
| F50874-06 | A | 21 | F | Gildersleeve |
| F50869-06 | AB | 30 | F | Gildersleeve |
| F50866-03 | B | 31 | M | Gildersleeve |
| F50869-09 | B | 30 | F | Gildersleeve |
| 541 | O | 44 | F | West |
| 655 | A | 34 | F | West |
| 663 | A | 34 | F | West |
| 688 | AB | 45 | M | West |
| 788 | B | NR | M | West |
| 791 | O | 33 | M | West |
| 794 | B | 41 | M | West |
| 796 | B | 68 | F | West |
| 826 | O | 28 | F | West |
| 844 | A | 27 | M | West |
| 1029 | O | 45 | F | West |
| 1109 | O | NR | M | West |
| 1114 | A | 69 | F | West |
| 1221 | O | 27 | M | West |
| 1293 | A | 51 | F | West |
| 1417 | O | NR | F | West |
| 1446 | O | 22 | F | West |
| 1471 | A | NA | M | West |

**Supplementary Table S1:** Information regarding serum samples used in this manuscript.

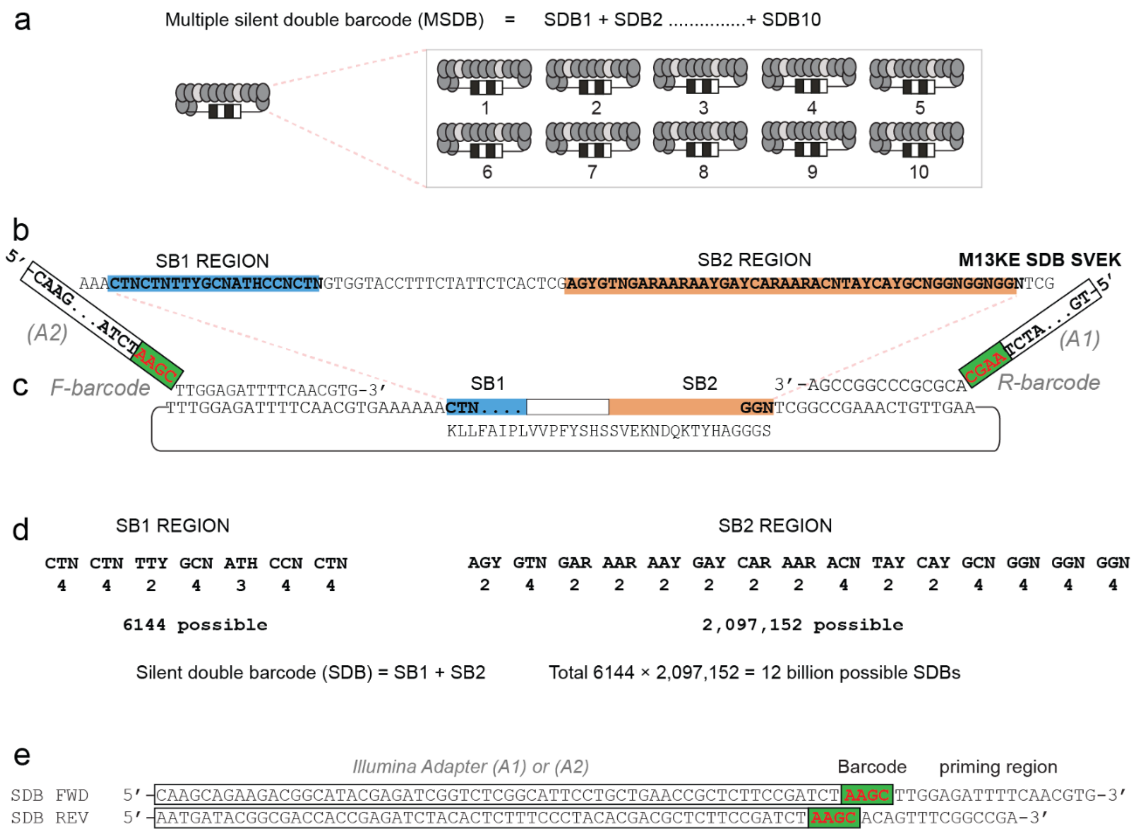

**Figure S1:** Scheme of silent double barcode (SDB) and multiple SDB (MSDB)
a) Illustration showing that MSDB is prepared by mixing individually cloned SDB stocks b) Each SDB clone contains two unique barcoding sites in M13-SDB vector termed silent barcode 1 and 2 (SB1 and SB2). These regions are located in pIII gene of M13 genome. c) Hybridization of Illumina seq primers to SDB vector d) Theoretical analysis of SDB diversity e, Sequences of the Illumina sequencing primers.

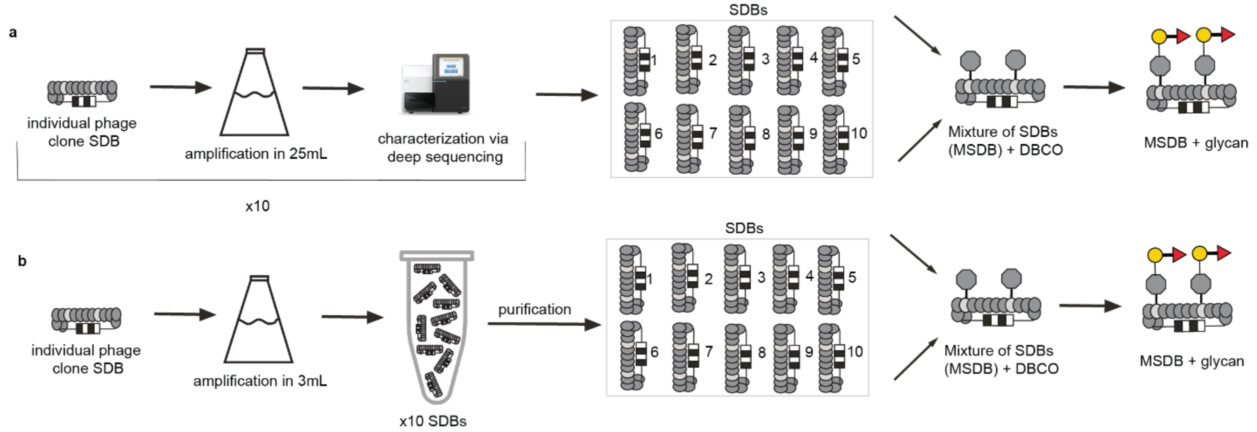

**Figure S2.** Construction of MSDBs.

a) Method 1 of constructing MSDBs. Individual phage clones were amplified in 25 mL flasks were characterized by deep sequencing before being mixed and conjugated to blood group antigens. b) Method 2 of constructing MSDBs. Individual phage clones were amplified separately in 3 mL volume and mixed together before purification of phage by PEG precipitation. Conjugation to blood group antigens followed.

| | B | C | D | E | |
|---|---|---|---|---|---|
| ...SDB | Position.... | Sequence........................................................ | Alphanum.. | # | Modification........... |
| SDB181 | [7:27 52:96] | TTATTATTCGCAATTCCTTTAAGTGTGGAGAAAAACGACCAAAAGACCTACCATGCTGGGGGGGGT | Tri-AN3-1 | 1 | Gala1-3[Fuca1-2]Galb-Sp |
| SDB12 | [7:27 52:96] | CTTCTGTTCGCGATACCTCTAAGTGTGGAGAAGAATGATCAGAAGACTTATCATGCGGGTGGAGGT | Tri-AN3-2 | 2 | Gala1-3[Fuca1-2]Galb-Sp |
| SDB16 | [7:27 52:96] | CTGCTGTTCGCAATCCCGCTGAGTGTGGAGAAGAATGATCAGAAGACTTATCATGCGGGTGGAGGT | Tri-AN3-3 | 3 | Gala1-3[Fuca1-2]Galb-Sp |
| SDB70 | [7:27 52:96] | CTACTGTTCGCAATCCCGCTCAGTGTTGAAAAAAACGATCAAAAAACGTATCATGCTGGTGGAGGT | Tri-AN3-4 | 4 | Gala1-3[Fuca1-2]Galb-Sp |
| SDB11 | [7:27 52:96] | CTACTATTTGCGATTCCCCTGAGTGTGGAGAAGAATGATCAGAAGACTTATCATGCGGGTGGAGGT | Tri-AN3-5 | 5 | Gala1-3[Fuca1-2]Galb-Sp |
| SDB25 | [7:27 52:96] | CTGCTTTTCGCAATACCTCTAAGTGTGGAGAAGAATGATCAGAAGACTTATCATGCGGGTGGAGGT | Tri-AN3-6 | 6 | Gala1-3[Fuca1-2]Galb-Sp |
| SDB45 | [7:27 52:96] | CTGCTGTTTGCGATTCCTCTGAGCGTGGAAAAAAATGACCAAAAAACCTACCATGCAGGGGGGGGA | Tri-AN3-7 | 7 | Gala1-3[Fuca1-2]Galb-Sp |
| SDB48 | [7:27 52:96] | TTATTATTCGCAATTCCTTTAAGCGTAGAGAAAAACGACCAGAAGACCTATCACGCGGGAGGAGGT | Tri-AN3-8 | 8 | Gala1-3[Fuca1-2]Galb-Sp |
| SDB49 | [7:27 52:96] | CTACTGTTCGCTATCCCGCTGAGTGTGGAAAAGAATGATCAGAAGACTTACCACGCTGGTGGTGGG | Tri-AN3-9 | 9 | Gala1-3[Fuca1-2]Galb-Sp |
| SDB50 | [7:27 52:96] | CTGCTCTTTGCGATTCCGCTGAGTGTCGAGAAGAACGACCAGAAAACATATCACGCGGGGGGGGGA | Tri-AN3-10 | 10 | Gala1-3[Fuca1-2]Galb-Sp |

**Figure S3:** Snapshot of dictionary that relates DNA sequences and glycan identities

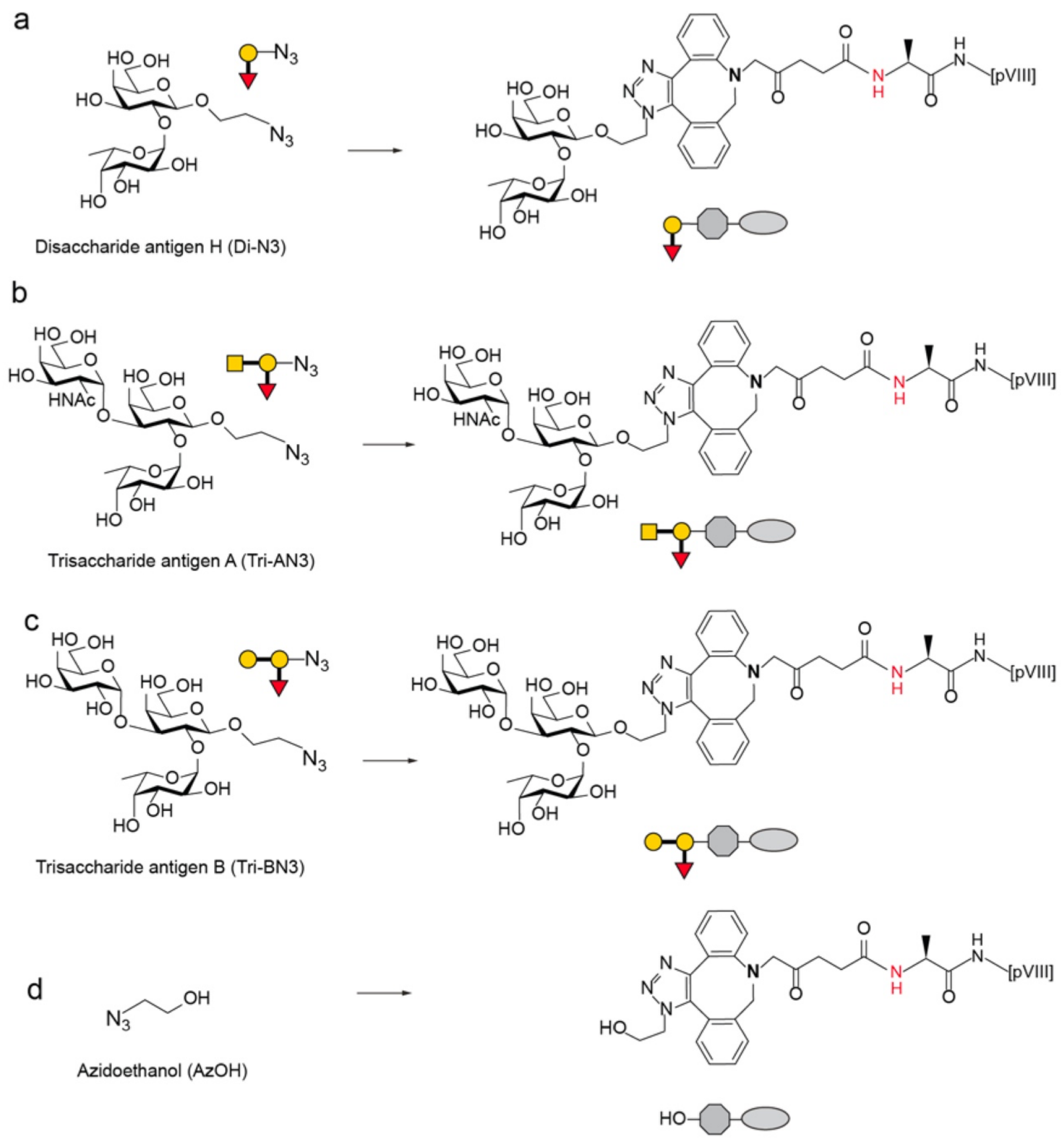

**Figure S4:** Structures of ABO glycans before and after conjugation to phage.
a) Disaccharide H b) Trisaccharide A c) Trisaccharide B d) Azidoethanol.

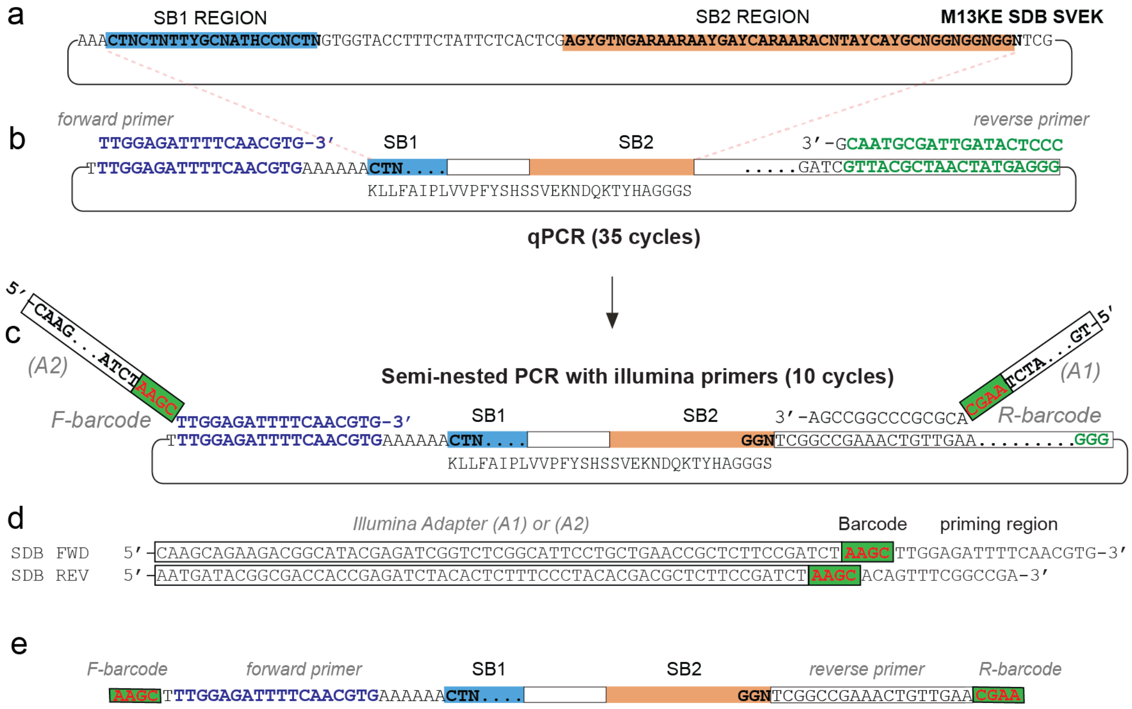

**Figure S2:**Scheme of qPCR and semi-nested PCR amplification.

a) Each SDB clone contains two unique sites in M13-SDB vector termed silent barcode (SB1) and (SB2 regions) b) first step qPCR with SDB forward primer without the Illumina overhangs and an outer reverse primer c) Second step semi-nested PCR with inner Illumina primers d) Sequences of the Illumina sequencing primers e) Region visible after Illumina sequencing of the second-step PCR product

a

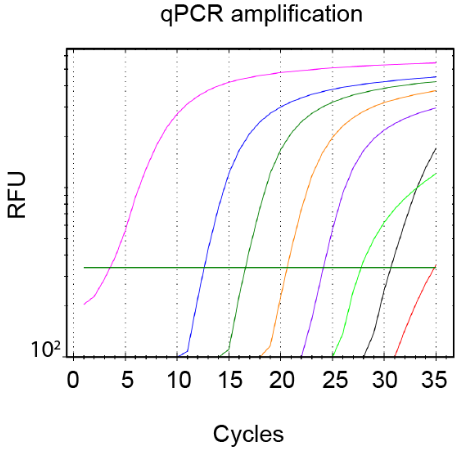

b

| Sample | PFU | Cq |
|---|---|---|
| | $10^8$ | 3 |
| | $10^7$ | 13 |
| | $10^6$ | 17 |
| | $10^5$ | 21 |
| | $10^4$ | 25 |
| | $10^3$ | 28 |
| | $10^2$ | 31 |
| | 10 | NA |
| | Neg | 35 |

**Figure S3:** Serial dilution-qPCR of an MSDB ABO LiGA library
a) qPCR amplification curve of different dilutions of a naïve MSDB library ranging from 10 to $10^8$ copies of phage calculated by PFU assay. b) Sample concentrations and their corresponding $C_q$ values

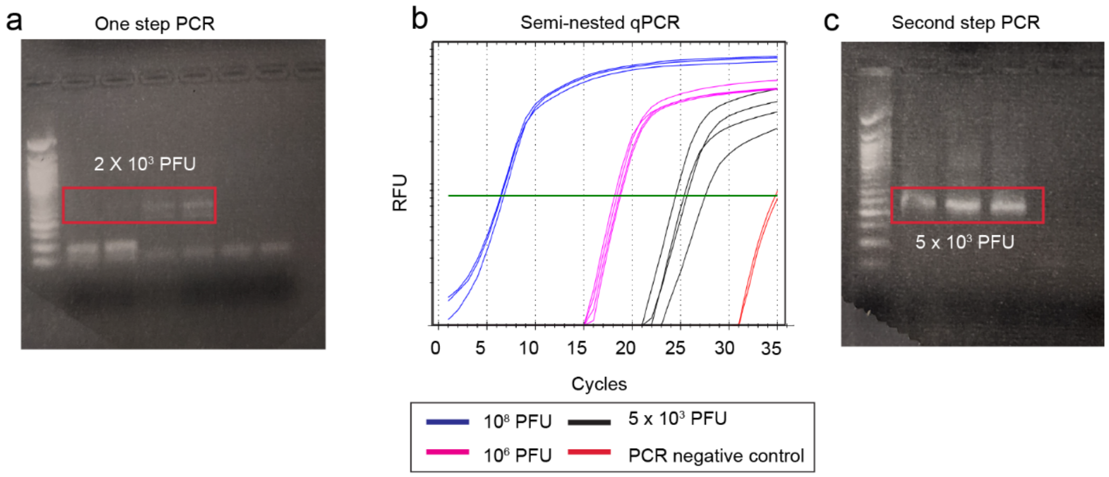

**Figure S4:** Comparison between one step and two step semi-nested PCR for low copy number phages

a) Gel electrophoresis image showing bands (highlighted in red) after one-step PCR amplification of phage with 2000 copies as PCR template b) qPCR amplification curve of phage samples recovered after a panning experiment c) Gel electrophoresis image showing bands after second-step semi nested PCR of low copy number phage with Illumina sequencing primers

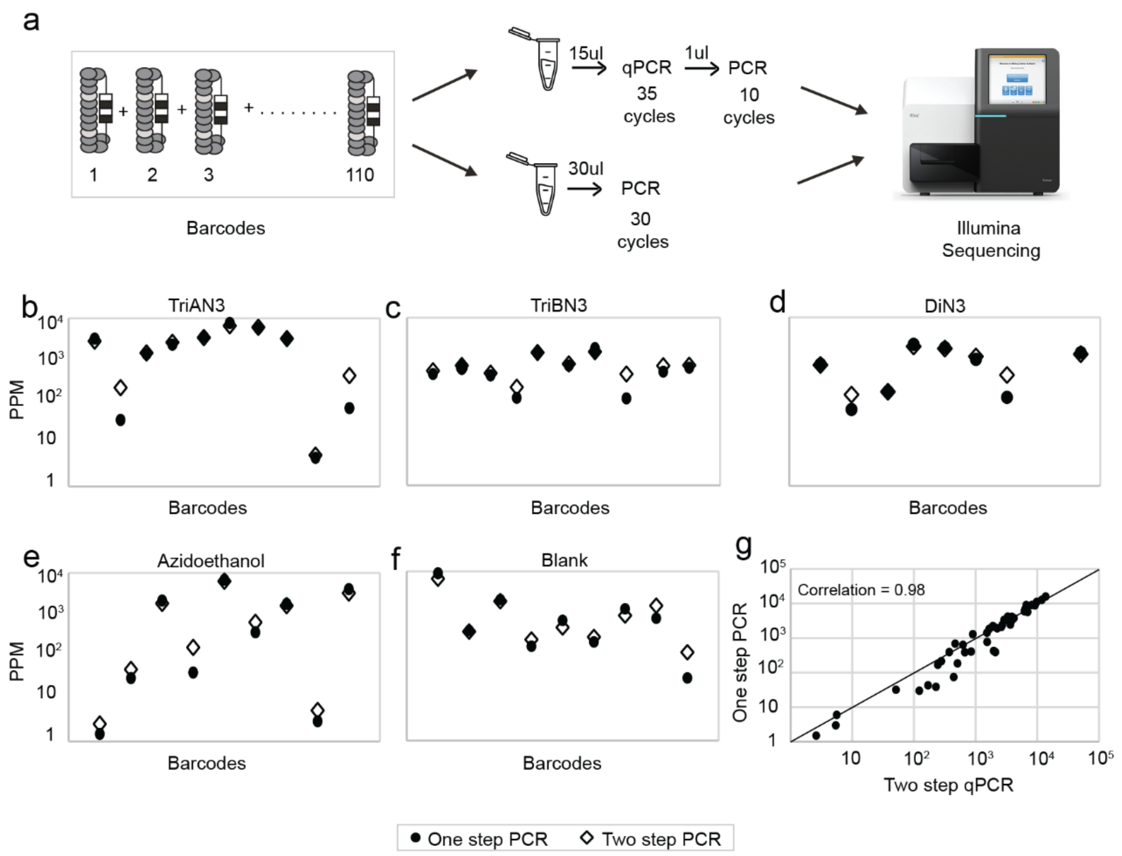

**Figure S5:** Comparison of Illumina deep sequencing results after one and two-step PCR. a) Illustration showing the sequencing workflow after PCR of a mixture consisting of 5 sets of MSDBs with $10^5$-$10^6$ templates used as input, each conjugated to either b) A blood group antigen (Tri-AN3), c) B blood group antigen (Tri-BN3), d) H(O) blood group antigen (Di-N3), or e) azidoethanol, f) unconjugated "blank" phages, g) MSDB ABO library PCR amplified using one step and two-step PCR showed little deviation of each clone's contribution to the total library sequenced with correlation coefficient of 0.98.

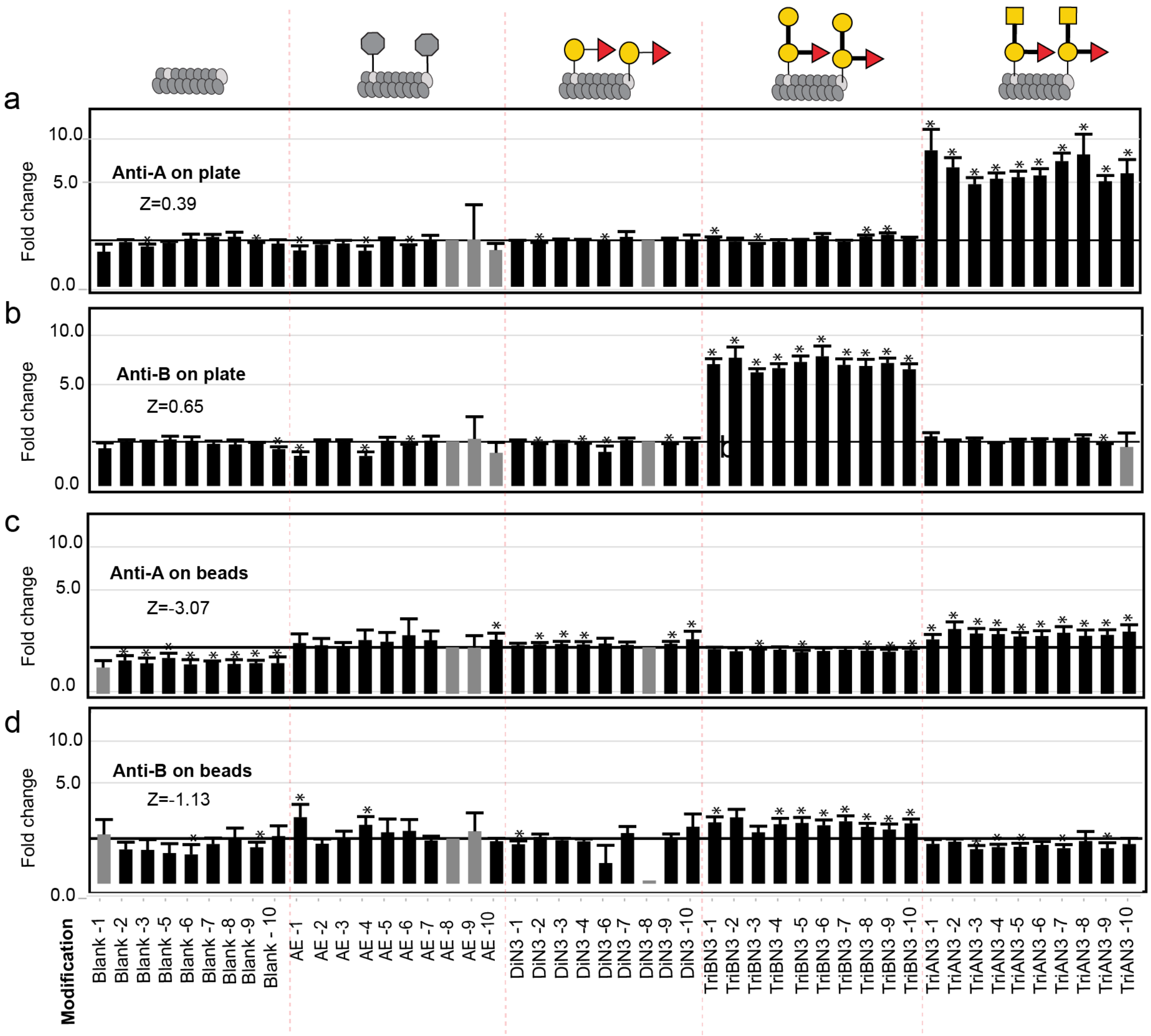

**Figure S6:** Deep sequencing analysis and Z' score calculation of ABO$_1$-LiGA with purified anti-A and anti-B IgM antibodies coated on **a,b,** plates and **c,d,** beads

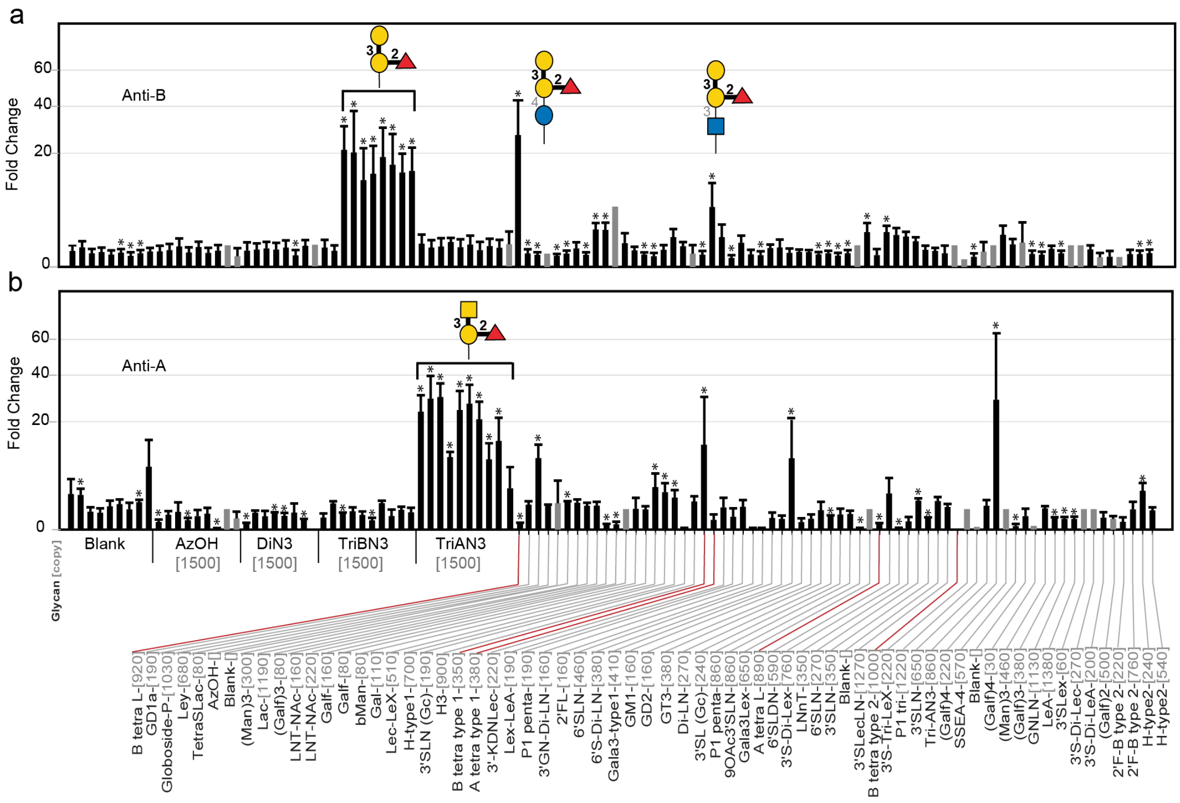

**Figure S7:** Screening ABO$_2$-LiGA library on anti-A and anti-B antibody.

ABO$_2$-LiGA was created by mixing ABO$_2$-LiGA with previously published LiGA library composed of 65 glycophage constructs (Sojitra et al. 2021) and incubated with plates coated by either anti-A or anti-B antibody. Red lines highlight ABO tetrasaccharide subtypes present in LiGA library and the table below has the IUPCA names for blood-group related glycans

| | |
|---|---|
| GalNAcα1-3[Fucα1-2]Galβ1-3GlcNAcβ-Sp | A tetra type 1 |
| GalNAcα1-3[Fucα1-2]Galβ1-4GlcNAcβ-Sp | A tetra type 2 |
| GalNAcα1-3[Fucα1-2]Galβ1-4Glcβ-Sp | A tetra L (A type VI) |
| GalNAcα1-3[Fucα1-2]Galβ1-4[Fucα1-3]GlcNAcβ-Sp | 2'F-A type 2 |
| Galα1-3[Fucα1-2]Galβ1-3GlcNAcβ-Sp | B tetra type 1 |
| Galα1-3[Fucα1-2]Galβ1-4GlcNAcβ-Sp | B tetra type 2 |
| Galα1-3[Fucα1-2]Galβ1-4Glcβ-Sp | B tetra L (B type VI) |
| Galα1-3[Fucα1-2]Galβ1-4[Fucα1-3]GlcNAcβ-Sp | 2'F-B type 2 |

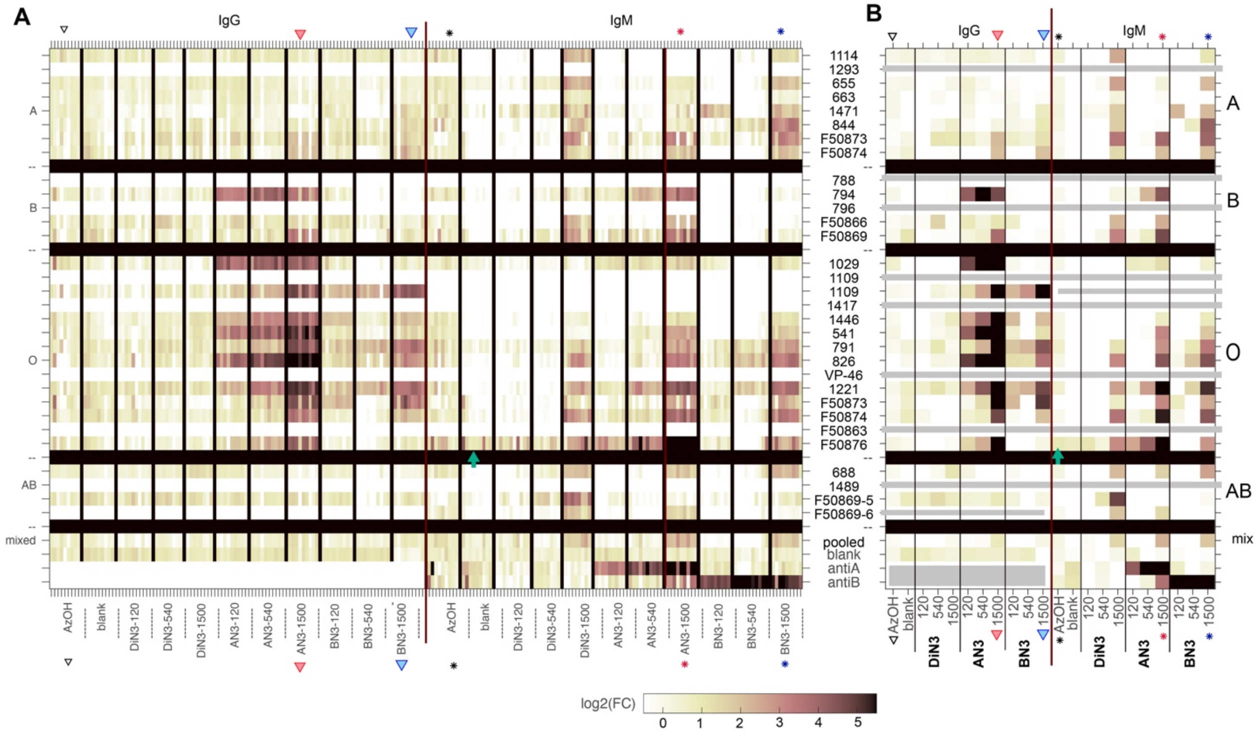

**Figure S11:** Breakdown of screening ABO LiGA described in Figure 4B to show the signal from every MSDB. Panel B is an exact copy from Figure 4B whereas panel A has the same layout as B but instead of showing the median signal, it shows the signal from every MSDB prior to the calculation of the median.

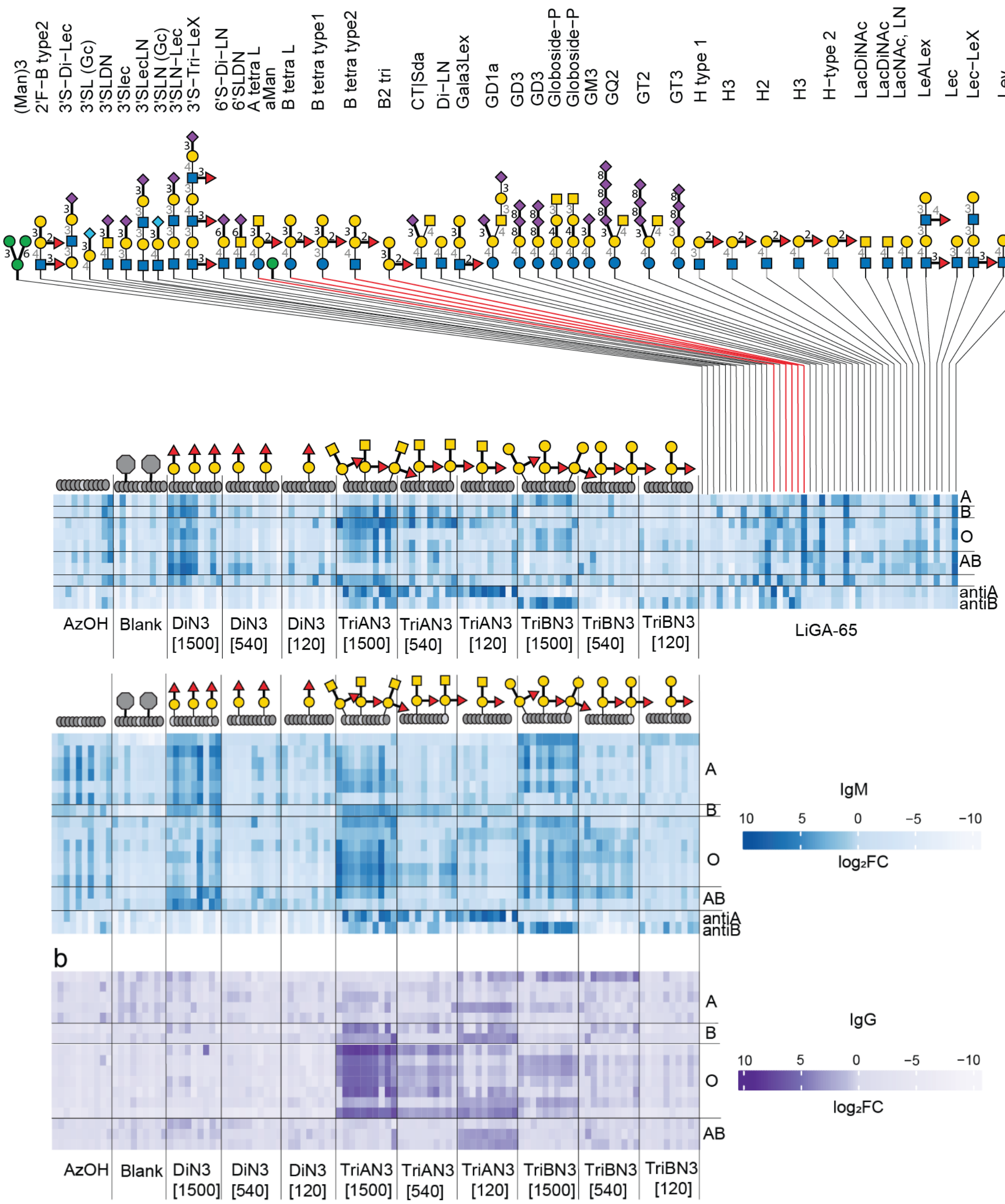

**Figure S12:** Heatmaps of sera IgM signals to ABO LiGA mixed with LiGA-65.
Subjects are clustered by blood-type in the rows. Glycans highlighted in red represent the ABO blood group glycans in the LiGA-65 mixture. Heatmap shows comparison of $log_2$ FC values for different glycans obtained by DE analysis. B) Heatmap describing the fold change enrichment of glycans in IgM and IgG-coated surface. Data is clustered by blood-type in the rows and by glycan density and glycan structure in the columns. For each density-structure, there are 10 DNA-encoded multi-SDB (MSDB) replicates and dispersion due to PCR amplification or NGS.

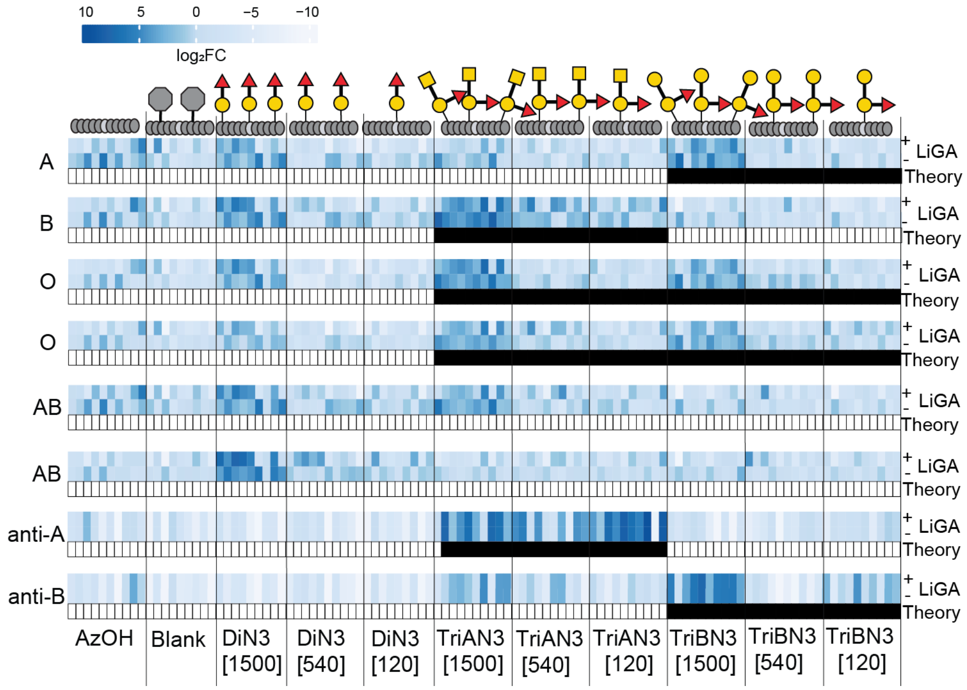

**Figure S13:** Comparison of IgM signals to $ABO_3$-LiGA with $ABO_4$-LiGA.

Both LIGA mixtures contain the same set of ABO-trisacharides displayed at three densities and $ABO_4$-LiGA contains additional glycans. For each serum samples, the top row is measured with $ABO_3$-LiGA and the bottom row is measured with $ABO_4$-LiGA. Black boxes illustrate an outcome of ABO-hemaglutination assay.

| Library | Sequencing file | Target |
|---|---|---|
| LiGA library EB ABO MSDBs | 20210305-87FAvgAW-RR | Blood group antibody A |
| | 20210305-87FAndAW-RR | Blood group antibody B |

**Supplementary Table S2:** Table of NGS raw data used in Figure 3.
The data may be accessed from https://48hd.cloud

| Blood type | Donor ID | Sequencing files for IgG | Sequencing files for IgM |
|---|---|---|---|
| A | 1114-1 | 20210219-87QLoiEG-RR | 20210219-87QLoiUF-RR |
| | 655-2 | 20210219-87QLdbEG-RR | 20210219-87QLdbUF-RR |
| | | 20210330-87FAdbEG-RR | 20210319-87FAtgUF-RR |
| | 663-1 | 20210219-87QLnhEG-RR | 20210219-87QLnhUF-RR |
| | | 20210330-87FAnhEG-RR | 20210330-87FAnhUF-RR |
| | 1471-1 | 20210330-87FAlcEG-RR | 20210330-87FAlcUF-RR |
| | 844-5 | 20210219-87QLjgEG-RR | 20210219-87QLjgUF-RR |
| | F50873-03 | 20210204-87QLufEG-RR | 20210204-87QLufUF-RR |
| | | 20210305-87FAufEG-RR | 20210305-87FAufUF-RR |
| | F50874-06 | 20210319-87FAieEG-RR | 20210319-87FAieUF-RR |
| B | 794-1 | 20210330-87FAldEG-RR | 20210330-87FAldUF-RR |
| | F50866-03 | 20210204-87QLwfEG-RR | 20210204-87QLwfUF-RR |
| | F50869-09 | 20210319-87FAhbEG-RR | 20210319-87FAhbUF-RR |
| O | 1029-7 | 20210330-87FAleEG-RR | 20210330-87FAleUF-RR |
| | 1109-5 | 20210219-87QLrhEG-RR | 20210219-87QLrhUF-RR |
| | 1446-1 | 20210219-87QLugEG-RR | 20210219-87QLugUF-RR |
| | | 20210330-87FAugEG-RR | 20210330-87FAugUF-RR |
| | 541-8 | 20210219-87QLkeEG-RR | 20210219-87QLkfUF-RR |
| | | 20210330-87FAkeEG-RR | 20210330-87FAkeUF-RR |
| | 791-1 | 20210219-87QLifEG-RR | 20210219-87QLifUF-RR |
| | 826-3 | 20210219-87QLfgEG-RR | 20210219-87QLfgUF-RR |
| | 1221-1 | 20210219-87QLrgEG-RR | 20210219-87QLrgUF-RR |
| | F50873-02 | 20210319-87FAeeEG-RR | 20210319-87FAeeUF-RR |
| | F50874-05 | 20210219-87QLkfEG-RR | 20210219-87QLkfUF-RR |
| | | 20210319-87FAkfEG-RR | 20210319-87FAkfUF-RR |
| | F50876-06 | / | 20210219-87QLniEG-RR |
| | | 20210319-87FAieEG-RR | 20210319-87FAieUF-RR |
| AB | 688-2 | 20210219-87QLohEG-RR | 20210219-87QLohUF-RR |
| | | 20210330-87FAohEG-RR | 20210330-87FAohUF-RR |
| | F50869-05 | 20210204-87QLtgEG-RR | 20210204-87QLtgUF-RR |
| | | 20210319-87FAtgEG-RR | 20210319-87FAtgUF-RR |
| | pooled | 20210219-87QLffEG-RR | 20210219-87QLffUF-RR |
| | blank | 20210305-87FAlfEG-RR | 20210305-87FAlgUF-RR |
| | antiA | | 20210305-87FAvgAW-RR |
| | antiB | | 20210305-87FAndAW-RR |

**Supplementary Table S3:** Table of NGS raw data used in Figure 4.
The data may be accessed from https://48hd.cloud

**Table S4: Summary of all NGS files collectred for this study.** Composition of LiGA QL, FA, are $ABO_3$-LiGA and $ABO_4$-LiGA described **Figure S12** and in QL.xlsx and FA.xsls as part of the supplementary files. All files can be downloaded from 48HD.cloud (search for the file name) or are available upon request.

| File Name | LiGA | Target |
|---|---|---|
| 20210204-87QLogUF-RR | QL | Human serum 869-06 coated on anti-human IgG |
| 20210204-87QLooPA-RR | QL | No target |
| 20210204-87QLtgEG-RR | QL | Human serum number 869-05 |
| 20210204-87QLtgUF-RR | QL | Human serum number 869-05 |
| 20210204-87QLufEG-RR | QL | Human serum number 873-03 |
| 20210204-87QLufUF-RR | QL | Human serum number 873-03 |
| 20210204-87QLwfEG-RR | QL | Human serum 866-03 |
| 20210204-87QLwfUF-RR | QL | Human serum 866-03 |
| 20210204-87QLwhMH-RR | QL | Protein L magnetic beads coated with anti-human IgM antibody |
| 20210219-87QLdbEG-RR | QL | Human serum 655-2_Lori West |
| 20210219-87QLdbUF-RR | QL | Human serum 655-2_Lori West |
| 20210219-87QLffEG-RR | QL | Human serum reference sample |
| 20210219-87QLffUF-RR | QL | Human serum reference sample |
| 20210219-87QLfgEG-RR | QL | Human serum 826_Lori West |
| 20210219-87QLfgUF-RR | QL | Human serum 826_Lori West |
| 20210219-87QLieUF-RR | QL | Human serum 874-06 |
| 20210219-87QLifEG-RR | QL | Human serum 791_Lori West |
| 20210219-87QLifUF-RR | QL | Human serum 791_Lori West |
| 20210219-87QLjgEG-RR | QL | Human serum 844_Lori West |
| 20210219-87QLjgUF-RR | QL | Human serum 844_Lori West |
| 20210219-87QLkeEG-RR | QL | Human serum 541_Lori west |
| 20210219-87QLkeUF-RR | QL | Human serum 541_Lori west |
| 20210219-87QLkfEG-RR | QL | Human serum 874-05_Lori West |
| 20210219-87QLkfUF-RR | QL | Human serum 874-05_Lori West |
| 20210219-87QLlfEG-RR | QL | Antihuman IgG coated on plates, used as a negative control for serum experiments |
| 20210219-87QLlgUF-RR | QL | Anti-human IgM coated on plates, used as blank/negative control for serum expts |
| 20210219-87QLnhEG-RR | QL | Human serum 663_Lori West lab |
| 20210219-87QLnhUF-RR | QL | Human serum 663_Lori West lab |
| 20210219-87QLniEG-RR | QL | Human serum 876-06_Lori West |
| 20210219-87QLohEG-RR | QL | Human serum 688_Lori West |
| 20210219-87QLohUF-RR | QL | Human serum 688_Lori West |
| 20210219-87QLoiEG-RR | QL | Human serum 1114-1_Lori West |
| 20210219-87QLoiUF-RR | QL | Human serum 1114-1_Lori West |
| 20210219-87QLooAW-RR | QL | No target |
| 20210219-87QLrgEG-RR | QL | Human serum 1221_Lori West |
| 20210219-87QLrgUF-RR | QL | Human serum 1221_Lori West |
| 20210219-87QLrhEG-RR | QL | Human serum 1109_Lori West |
| 20210219-87QLrhUF-RR | QL | Human serum 1109_Lori West |

| | | |
|---|---|---|
| 20210219-87QLugEG-RR | QL | Human serum 1446_Lori West |
| 20210219-87QLugUF-RR | QL | Human serum 1446_Lori West |
| 20210305-87FAbwAW-RR | FA | Blank well |
| 20210305-87FAlfEG-RR | FA | Antihuman IgG coated on plates, used as a negative control for serum experiments |
| 20210305-87FAlgUF-RR | FA | Anti-human IgM coated on plates, used as blank/negative control for serum expts |
| 20210305-87FAndAW-RR | FA | Blood group antibody B |
| 20210305-87FAooPA-RR | FA | No target |
| 20210305-87FAufEG-RR | FA | Human serum number 873-03 |
| 20210305-87FAufUF-RR | FA | Human serum number 873-03 |
| 20210305-87FAvgAW-RR | FA | Blood group antibody A |
| 20210305-87FAwfEG-RR | FA | Human serum 866-03 |
| 20210305-87FAwfUF-RR | FA | Human serum 866-03 |
| 20210305-87KCwfUF-RR | KC | Human serum 866-03 |
| 20210305-87FAbwAW-RR | FA | Blank well |
| 20210319-87FAeeEG-RR | FA | Human serum number 873-02 |
| 20210319-87FAeeUF-RR | FA | Human serum number 873-02 |
| 20210319-87FAffUF-RR | FA | Human serum reference sample |
| 20210319-87FAhbEG-RR | FA | Human serum 869-09 |
| 20210319-87FAhbUF-RR | FA | Human serum 869-09 |
| 20210319-87FAieEG-RR | FA | Human serum 874-06 |
| 20210319-87FAieUF-RR | FA | Human serum 874-06 |
| 20210319-87FAkfEG-RR | FA | Human serum 874-05_Lori West |
| 20210319-87FAkfUF-RR | FA | Human serum 874-05_Lori West |
| 20210305-87FAlfEG-RR | FA | Antihuman IgG coated on plates, used as a negative control for serum experiments |
| 20210305-87FAlgUF-RR | FA | Anti-human IgM coated on plates, used as blank/negative control for serum expts |
| 20210305-87FAndAW-RR | FA | Blood group antibody B |
| 20210319-87FAniEG-RR | FA | Human serum 876-06_Lori West |
| 20210319-87FAniUF-RR | FA | Human serum 876-06_Lori West |
| 20210319-87FAogUF-RR | FA | Human serum 869-06 coated on anti-human IgG |
| 20210305-87FAooPA-RR | FA | No target |
| 20210319-87FAtgEG-RR | FA | Human serum number 869-05 |
| 20210319-87FAtgUF-RR | FA | Human serum number 869-05 |
| 20210305-87FAufUF-RR | FA | Human serum number 873-03 |
| 20210305-87FAvgAW-RR | FA | Blood group antibody A |
| 20210305-87FAbwAW-RR | FA | Blank well |
| 20210330-87FAdbEG-RR | FA | Human serum 655-2_Lori West |
| 20210330-87FAdbUF-RR | FA | Human serum 655-2_Lori West |
| 20210330-87FAkeEG-RR | FA | Human serum 541_Lori west |
| 20210330-87FAkeUF-RR | FA | Human serum 541_Lori west |
| 20210330-87FAlcEG-RR | FA | Human serum sample 1471 in heparin, blood group A, Lori West group |
| 20210330-87FAlcUF-RR | FA | Human serum sample 1471 in heparin, blood group A, Lori West group |
| 20210330-87FAldEG-RR | FA | Human serum sample 794 in heparin, blood group B, Lori West group |
| 20210330-87FAldUF-RR | FA | Human serum sample 794 in heparin, blood group B, Lori West group |
| 20210330-87FAleEG-RR | FA | Human serum sample 1029, serum, Blood group O, Lori West group |
| 20210330-87FAleUF-RR | FA | Human serum sample 1029, serum, Blood group O, Lori West group |

| | | |
|---|---|---|
| 20210305-87FAlfEG-RR | FA | Antihuman IgG coated on plates, used as a negative control for serum experiments |
| 20210305-87FAlgUF-RR | FA | Anti-human IgM coated on plates, used as blank/negative control for serum expts |
| 20210305-87FAndAW-RR | FA | Blood group antibody B |
| 20210330-87FAnhEG-RR | FA | Human serum 663_Lori West lab |
| 20210330-87FAnhUF-RR | FA | Human serum 663_Lori West lab |
| 20210330-87FAohEG-RR | FA | Human serum 688_Lori West |
| 20210330-87FAohUF-RR | FA | Human serum 688_Lori West |
| 20210305-87FAooPA-RR | FA | No target |
| 20210330-87FAugEG-RR | FA | Human serum 1446_Lori West |
| 20210330-87FAugUF-RR | FA | Human serum 1446_Lori West |
| 20210305-87FAvgAW-RR | FA | Blood group antibody A |
